# ARACoFusion: Uncertainty-aware calibrated deep learning for protein-protein interaction network prediction in *Arabidopsis thaliana*

**DOI:** 10.64898/2026.05.22.727120

**Authors:** Dipayan Sarkar, Chiranjib Sarkar

## Abstract

Accurate mapping of the *Arabidopsis thaliana* protein-protein interaction (PPI) network is essential for deciphering complexity of plant systems biology. Here, we present ARACoFusion, a specialized deep learning architecture designed to predict inter-protein connectivity directly from primary sequences. To capture the asymmetric dependencies between plant proteins, the framework utilizes a reciprocal cross-attention encoder combined with latent interaction projections and multi-source feature fusion. Addressing the severe class imbalance inherent in plant interactomes, the model integrates uncertainty-aware variance regularization and focal loss with label smoothing, further enhancing reliability through post-hoc probability calibration via temperature scaling. Extensive benchmarking on gold-standard Arabidopsis datasets demonstrates that ARACoFusion significantly outperforms existing plant-specific predictors, achieving superior scores in Area Under the Precision-Recall Curve (AUPRC), Balanced Accuracy, and Matthew’s Correlation Coefficient (MCC). Additionally, the model exhibits robust cross-species generalization and clear class separability in t-SNE latent space visualizations. To facilitate community-wide usage, we provide a dedicated web server for scalable network-level inference at https://ARAcofusion.compbiosysnbu.in/.

## 1 Introduction

From developmental regulation to environmental stress adaptation, the physiological plasticity of *Arabidopsis thaliana* is fundamentally coordinated by dynamic protein-protein interaction (PPI) networks. These molecular interactomes act as the cellular backbone, driving signal transduction, metabolic channeling, and hormone cross-talk. System-level analyses have demonstrated that one cannot decipher complex traits such as defense metabolite biosynthesis by studying isolated proteins. Instead, a holistic mapping of the network architecture is required [1,2]. To reconstruct these networks, high-throughput experimental assays including yeast two-hybrid screens [3], tandem affinity purification [4], and mass spectrometry [5] have been instrumental in generating initial interaction skeletons. However, these wet-lab approaches face significant scalability challenges when applied to the entire plant proteome. They are often hindered by high rates of stochastic noise, creating datasets prone to both false positives (spurious interactions) and false negatives (missed transient interactions). Consequently, a vast portion of the *Arabidopsis* interactome remains unmapped, necessitating computational interventions to bridge the gap between genomic data availability and functional network knowledge. Existing computational strategies for PPI prediction are generally categorized into docking-based [6], structure-based [7], and sequence-based approaches [8]. While structural methods offer atomic-level resolution, they are severely constrained by the scarcity of experimentally solved 3D structures for plant proteins. In contrast, sequence-based methodologies leverage the exponential growth of genomic data, learning predictive patterns directly from primary amino acid chains without requiring prior structural annotation. Historically, machine learning efforts in this domain relied on handcrafted feature engineering to translate biological sequences into numerical formats. Techniques such as amino-acid composition (AAC), conjoint-triad (CT), and dipeptide composition (DPC) were frequently employed to extract physicochemical descriptors. These static features served as inputs for classical classifiers, including Support Vector Machines (SVM) [9], Random Forest [10], and K-Nearest Neighbor algorithms [11]. While these methods established the feasibility of in silico prediction, they utilize static representations that often fail to capture the complex, non-local dependencies and evolutionary context essential for high-accuracy interaction inference.

In recent years, deep learning has emerged as the dominant paradigm for sequencebased PPI prediction [12], driven by its ability to extract high-dimensional features and model non-linear dependencies via back-propagation. Within this domain, Natural Language Processing (NLP) techniques have proven to be a biologically intuitive approach for protein embedding. Initial applications utilized static embedding techniques such as Word2Vec or Doc2Vec, which treat amino acids and peptide segments analogously to words in a sentence [13]. While these unsupervised methods offered an improvement over manual feature engineering, they remain “shallow”, struggling to capture long-range dependencies or the complex hierarchical Patterns of protein sequences. To address these shortcomings, Transformer-based architectures have revolutionized sequence modeling through self-attention mechanisms [14], which capture global context regardless of sequence length. Pretrained protein language models (PLMs) such as ProtBERT [15], ESM-1b [16], and ProteinBERT [17] have since achieved state-of-the-art performance. By distilling evolutionary patterns from millions of unlabelled sequences, these models significantly enhance the accuracy and generalization of downstream PPI predictors [18]. However, general-purpose models trained on multi-species data often fail to capture the unique evolutionary signals and extreme class imbalance inherent to specific plant interactomes, particularly in *Arabidopsis thaliana*.Efforts to develop *Arabidopsis* specific predictors have yielded tools like AraPPINet [19], which offers insights into hormone signaling pathways by combining genomic context with interolog mapping. However, AraPPINet relies heavily on structure-derived features, severely limiting its applicability to the vast majority of plant proteins that lack solved 3D structures. Subsequent approaches, such as DeepAraPPI [20], attempted to bridge this gap by integrating Word2Vec and GO2Vec embeddings into a logistic regression framework. Although this improved upon traditional machine learning baselines, the model remains dependent on handcrafted features and focuses predominantly on pairwise classification rather than holistic network modeling.More recently, the ESMAraPPI framework [21] advanced the field by applying the ESM-1b language model to encode sequence semantics, utilizing a Multilayer Perceptron (MLP) for binary classification. While effective, this architecture is limited by its simplicity; it does not account for the probabilistic uncertainty inherent in biological data, nor does it address the significant generalization gap observed when models are deployed on highly unbalanced, real-world datasets.

To overcome these methodological constraints, we introduce ARACoFusion, a specialized deep learning architecture designed for robust PPI prediction in *Arabidopsis*. Unlike previous models, ARACoFusion moves beyond simple binary classification by integrating a reciprocal cross-attention encoder with latent interaction projection and multi-source feature fusion. Recognizing the challenge of noise in high-throughput datasets, we incorporate uncertainty-aware training via variance regularization to stabilize predictions and reduce overfitting. Furthermore, we apply temperature scaling as a post-hoc calibration mechanism to ensure that predicted probabilities reflect true interaction likelihoods. The entire architecture is fine-tuned using Bayesian hyperparameter optimization via Optuna [22]. By combining these advanced components, ARACoFusion provides a highly generalizable, calibration-aware framework capable of mapping high-confidence interaction subnetworks within the complex landscape of plant systems biology.

## 2 Materials and Methods

### 2.1 Datasets

The experimentally validated gold standard *Arabidopsis thaliana* datasets were downloaded from IntAct (https://www.ebi.ac.uk/intact/home), obtained from Zhou et al. [21]. The dataset having interaction pairs with physical association and MI score of < 0.45 was retained, constructing the final dataset of 7729 positive samples. The negative dataset was constructed using three steps. First, the positive samples obtained earlier were removed from the complete dataset, and the remaining protein sequences having > 40% sequence identity with positive samples were also removed. Secondly, the proteins with similar sequence identity were removed using a 40% cutoff, and 8382 proteins were obtained. Thirdly, the final negative protein pair list was made using those 8382 proteins and the proteins in positive samples by random pairing. The final dataset was constructed with a 1:10 ratio of positive and negative samples, comprising 7729 positive and 77290 negative samples. To construct a training and evaluation dataset, the final data were divided into three datasets as C1, C2, and C3 using Park and Marcotte’s method [23]. C1 is used for training, and C2 and C3 are used for validation. The data distribution is visible in Table1. For cross-species validation experimentally verified rice PPI dataset downloaded from four public databases (DIP, MINT, BioGRID, and IntAct) was obtained from Zheng et al. [20]. Self, non-physical, and redundant interactions were removed to obtain 611 positive interactions. Protein pairs that were not rice PPIs were randomly selected as positive samples, obtaining the final dataset with a 1:10 ratio of 611 positive and 6111 negative interactions.

**Table 1.**
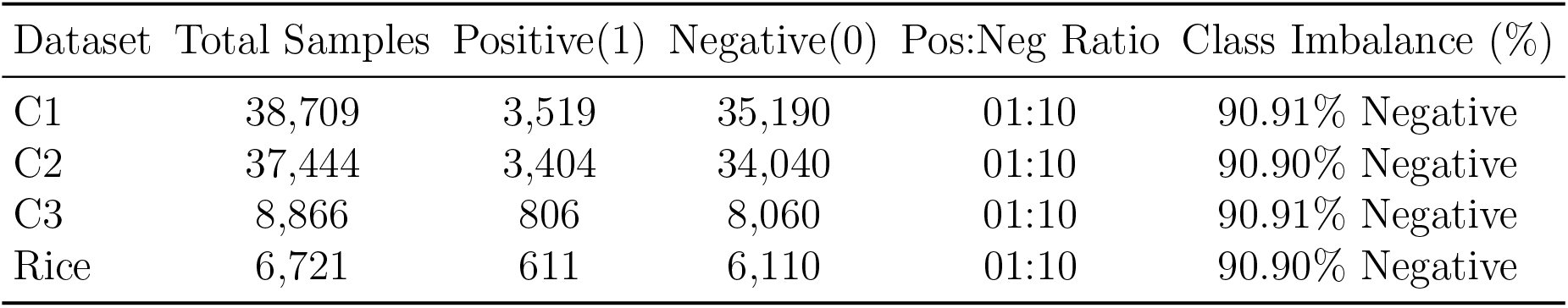
Data distribution and class imbalance.

| Dataset | Total Samples | Positive(1) | Negative(0) | Pos:Neg Ratio | Class Imbalance (%) |
| --- | --- | --- | --- | --- | --- |
| C1 | 38,709 | 3,519 | 35,190 | 01:10 | 90.91% Negative |
| C2 | 37,444 | 3,404 | 34,040 | 01:10 | 90.90% Negative |
| C3 | 8,866 | 806 | 8,060 | 01:10 | 90.91% Negative |
| Rice | 6,721 | 611 | 6,110 | 01:10 | 90.90% Negative |

### 2.2 Protein encoding method

In the post-Transformer era, with the advent of attention mechanisms [14], protein sequence embedding revolutionised from hand-crafted or unsupervised features to highdimensional structure-guided, evolution-scaled, feature-rich Protein language models (PLMs) trained on billions of proteins. Unlike our previous framework [18], which relied on ProtT5-XL embeddings, ARACoFusion employs the the ESM 1b-650M [16] parameter model developed by Meta AI, which converts raw amino-acid sequences into a highdimensional contextual embedding.We found that ESM-1b’s evolutionary-scale training provides superior resolution for capturing the subtle co-evolutionary signals required for plant-specific interaction prediction. Using a masked language model objective, ESM1 was pretrained with the UniRef50 dataset, comprising 40 million proteins. Unlike earlier language models like TAPE or UniRep, ESM 1b captures long-range dependencies and structural features only from sequence data, outperforming its predecessor in several benchmarks. Depending on the parameters, several versions of ESM models are available, ranging from 8 million to 15 billion parameters, and depending on that, the embedding dimension also increases or decreases. The raw embeddings were generated for each amino acid and were concatenated to produce a mean embedding representing each protein (Figure 1).If a single protein sequence *S* = (*s*_1_, *s*_2_, *s*_3_, ……, *s*_*L*_) where the protein sequence length is *L* and contextual embedding *x* of *i*-th resedue from the final layer of ESM 1b 650M model with *dimension*(*d*) = 1280, is *x*^*i*^ ∈ ℝ;^*d*^, then the per protein embedding *z* ∈ ℝ^*d*^ is

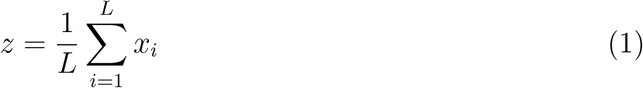

**Figure 1.**
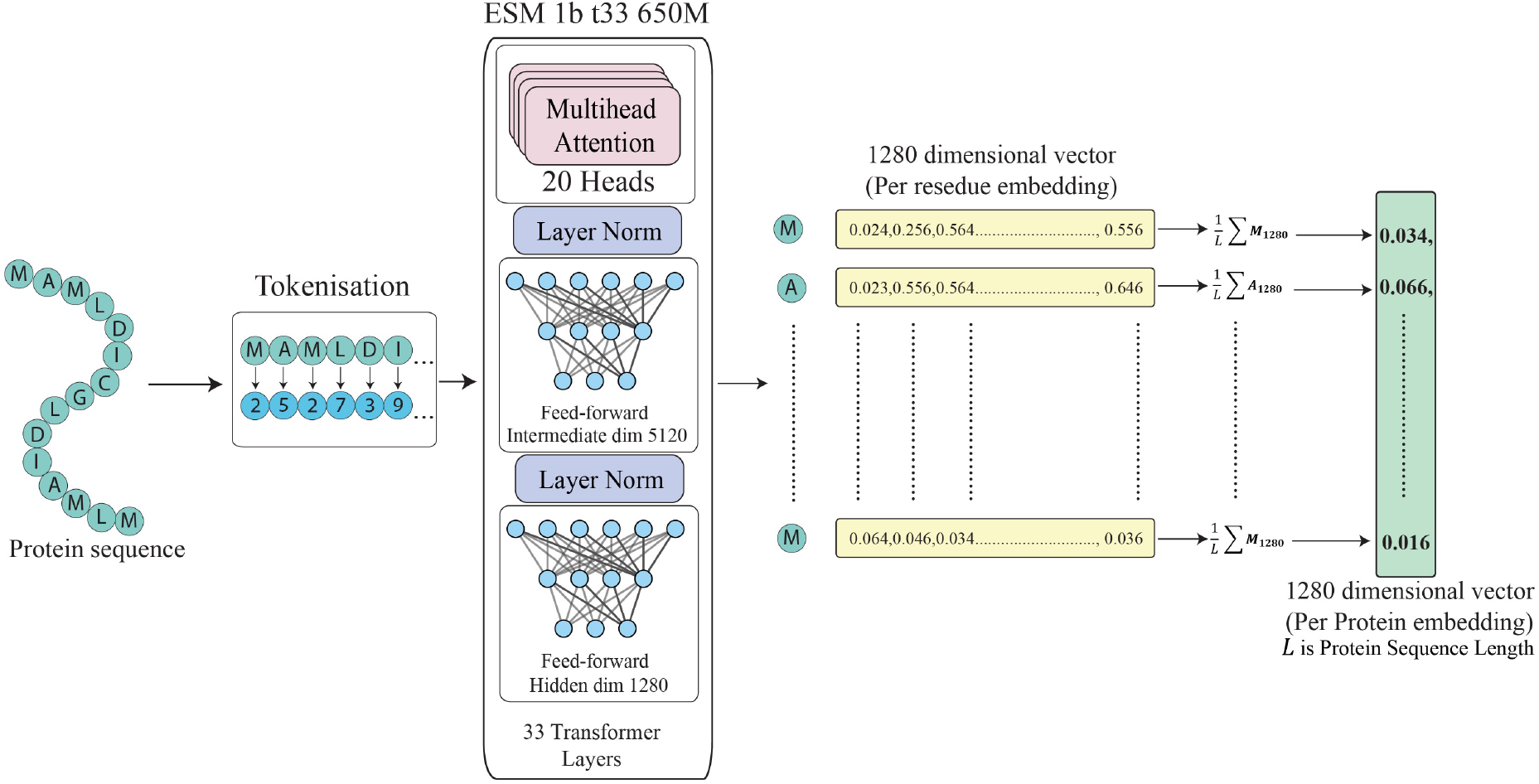
Protein embedding generation using ESM-1b T33 650M. Protein sequences are tokenized and processed through 33 transformer layers to produce 1280-dimensional per-residue embeddings. These are averaged across the sequence length to obtain a fixed-size 1280-dimensional per-protein embedding.

### 2.3 ARACoFusion model architecture

We developed a novel deep neural network architecture named ARACoFusion to predict protein-protein interaction directly from per-protein sequence embeddings (Figure 2). This architecture has three primary components: a reciprocal cross-attention Encoder, a latent interaction projector, and a multi-stage fusion-based classifier head.

**Figure 2.**
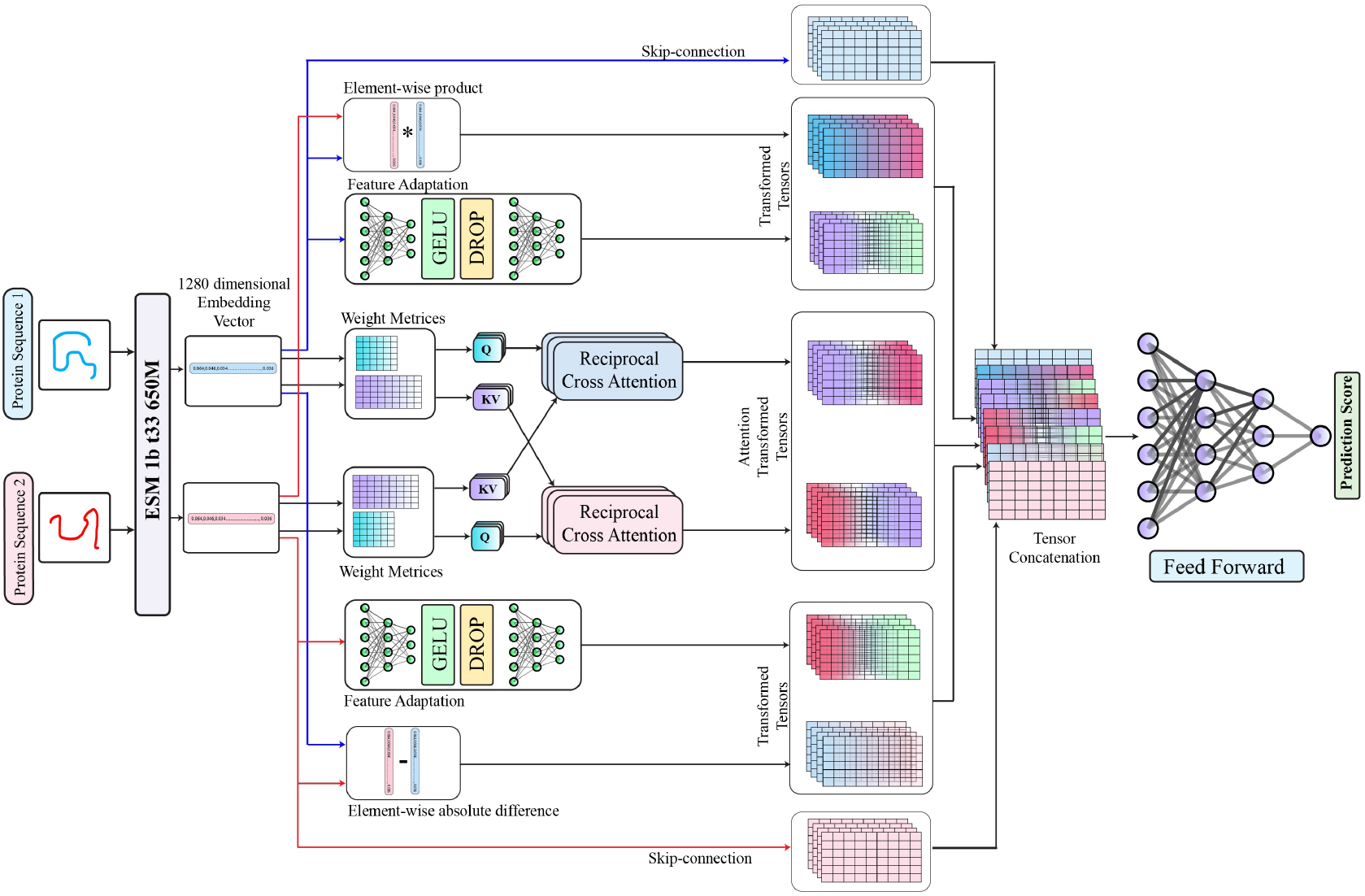
Architecture of the ARACoFusion model for Arabidopsis PPI prediction. Protein sequences are encoded using the ESM 1b-t33-650M model to generate 1280-dimensional embeddings. Each embedding passes through reciprocal cross-attention layers, where one protein attends to the other via learned query-key-value transformations. Latent interaction features are further refined through nonlinear feature adaptation modules and enriched by element-wise product and absolute difference operations. All transformed features are concatenated and passed through a multi-layer feedforward network to generate the final interaction probability. Skip-connections preserve original context, enabling stable gradient flow and robust fusion.

#### 2.3.1 Input representation

The input of architecture is a pair of protein sequences and each embedded in a fixed-length vector. The language model ESM-1b-650M embeds each protein sequence into a 1280-dimensional (*d*) vector.The input vectors are concatenated to form tensors and divided into batches for batch processing. If the input tensor *X* ∈ ℝ^*B*×2×*d*^ is decomposed as

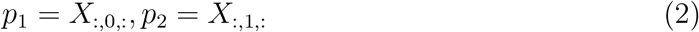

where, *B* is the batch size, *p*_1_ and *p*_2_ is batched tensor of protein1 and protein2 consecutively.

#### 2.3.2 Reciprocal Cross-attention Encoder

The architecture employs a reciprocal multi-head cross-attention encoder, which enables each protein embedding to selectively attend to context from other protein embeddings. Linear transformations are applied to generate queries (*Q*), key-values (*KV*) for each protein embedding. If *p*_1_*p*_2_ ∈ ℝ^*d*^, where the embedding dimension *d* = 1280, then the protein embedding *p*_1_ : *Q*_1_ = *W*_*q1*_ *p*_1_, where *Q*_1_ ∈ ℝ^*d*^ and 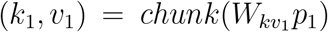, where *K*_1_, *V*_1_ *∈* ℝ^*d*^ and for protein embedding *p*_2_ : *Q*_2_ = *W*_*q*2_ *p*_2_, where *Q*_2_ ∈ ℝ and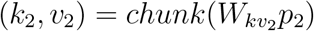, where *k*_2_, *V*_2_ ∈ ℝ^*d*^.

Here *W*_*q i*_ *∈* ℝ^*d*×*d*^, and 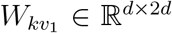, which represent learnable weight metrics. The *chunk* operation splits the output tensor evenly into key and value tensors, each with dimension *d*. Each protein embedding reciprocally attends to key-value pairs of other protein embeddings. When protein 1 attends to protein 2, to compute the attention scores and contextual embedding (*c*_1_) for *p*_1_:

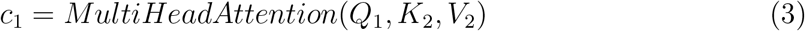

where, the multi head attention with H heads, computed by

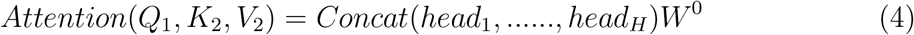

Each head can be defined as

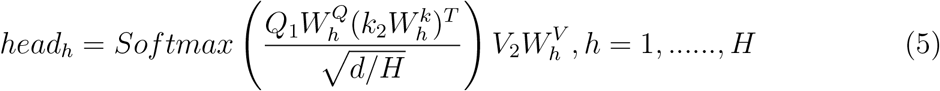

Here, 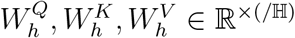 are learned projection metrics for each head *h*.*W* ^0^ *∈* ℝ^*d×d*^ which combines the output from all attention heads. Similarly, when protein 2 attends to protein 1, to compute the attention scores and contextual embedding (*c*_2_) for *p*_2_.

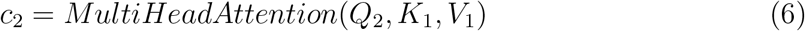

The *c*_1_ and *c*_2_ represents cross-contextualised embeddings for each protein, where *c*_1_, *c*_2_∈ ℝ^*d*^.

#### 2.3.3 Feature adaptation and Latent interaction projector

In the feature adaptation state, two feature enhancement strategies were adopted, when the two protein embeddings *p*_1_, *p*_2_ ∈ ℝ^*d*^ with *d* = 1280. We derived explicit pairwise relational features as element-wise product and element-wise absolute difference. Element-wise product (*P*_*p*_*rod*)

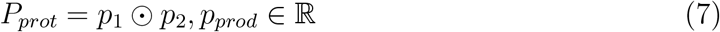

This captures the direct pairwise compatibility and co-activating features between corresponding embeddings.Element-wise absolute difference *P*_*diff*_

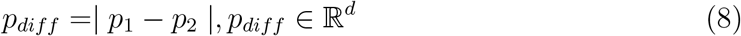

This measures the divergence between embedding, indicating regions of substantial embedding difference. In parallel, each protein embedding undergoes a nonlinear adaptation through a two-layer fully connected neural network, the goal is to capture deeper, latent representation and to reduce embedding dimensionality. For protein embedding *p*_*i*_, the latent interaction can be projected as

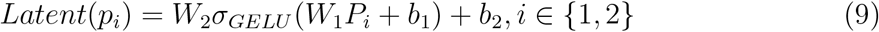

Where, *W*_1_, *W*_2_, *b*_1_, *b*_2_ are weights and biases of the linear transformation layer 1 and 2, respectively, and *σ*_*GELU*_ denotes the Gaussian Error Linear Unit activation function. Thus, each latent represents an embedding *a*_*i*_ is

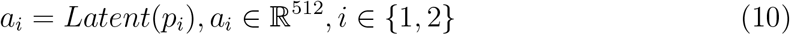

#### 2.3.4 Concatenation and multi-stage fusion-based classifier head

To form the final interaction representation (*Z*), we concatenated the original embeddings *p*_1_, *p*_2_; reciprocal cross-attention embeddings *c*_1_, *c*_2_; element-wise interactions *P*_*prod*_, *P*_*diff*_ ; and latent representations *a*_1_, *a*_2_. Thus, the final comprehensive interaction representation is

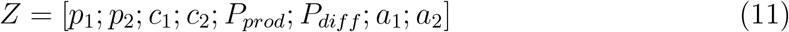

The interaction prediction head comprises multiple fully connected layers, which can be represented as

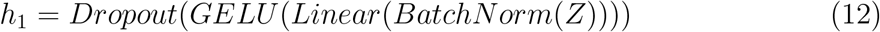

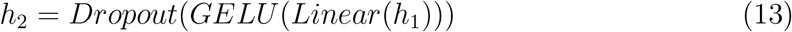

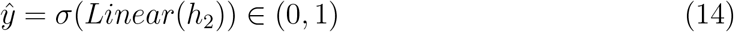

where, the *σ* is the sigmoid operation, i.e. 1*/*(1+*e*^−*z*^) to produce an interaction probability score.

### 2.4 Model Regularization, Calibration, and Optimization

#### 2.4.1 Uncertainty-aware Training with Variance Regularization

The model was incorporated with a form of uncertainty estimation to improve its generalization ability and to reduce over-fitting. The uncertainty was estimated through a variance-based regularization term in the loss function, first introduced by Gal et al. [24]. During each training step, the model performs Monte Carlo sampling by repeating the forward pass three times for the same input batch. For each input pair, we computed the variance of the predicted probabilities across these passes. The high variance indicates uncertainty about the model’s prediction. We added a penalty to the training loss proportional to this variance, which encourages the model to make more confident and stable predictions. The final training loss can be defined as

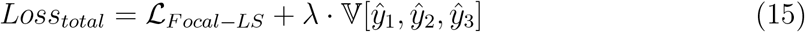

Here, *L*_*F ocal−LS*_ represents the focal loss with label smoothing, which is used to handle class imbalance. 𝕍 is the variance of predictions *ŷ*_1_, *ŷ*_2_, *ŷ*_3_ obtained from these three passes. The confidence *λ* controls the strength of the variance penalty, which is set using hyperparameter optimization.

#### 2.4.2 Temperature scaling for probability calibration

Temperature scaling [25] was applied to reduce overconfident prediction and to improve the reliability of neural networks. Temperature scaling is a post-training calibration technique that adjusts the model’s confidence without changing its classification accuracy. A single scaler parameter called the temperature (*T*), which was optimized using the logits from the validation set, is used to rescale the output logits. The rescaled logits then passed through a sigmoid function to compute calibrated probabilities

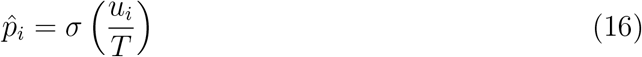

where, *u*_*i*_ is the uncalibrated logits for the *i*-th sample, *σ*(·) is the sigmoid function, 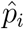 is the calibrated probability, *T >* 0 is the temperature parameter.

#### 2.4.3 Hyperparameter optimization using Optuna

We employed automated hyperparameter optimization using Optuna [22] to enhance the performance and generalization of the model. Optuna is an efficient and flexible hyper-parameter search library that uses the Tree-structure Parzen Estimator (TPE) algorithm to guide the optimization process. In our study Optuna was used to find the optimal values of the hyperparameter, including learning-rate, dropout-rate, focal-loss gamma, and smoothing factor, and *λ* parameter in uncertainty-aware variance regularization.

### 2.5 Model validation techniques

The proposed model was evaluated using C1 as train, C2, C3 as validation data with 5-fold cross validation, cross species prediction and PPI network prediction. To ensure robust performance evaluation, we employed stratified 5-fold cross-validation. This procedure partitions the dataset into five distinct subsets while preserving the ratio of interacting to non-interacting pairs in each fold, thereby preventing bias during training and testing. For validation of ARACoFusion model, the following statistical measures were used to evaluate the performance of the trained model:

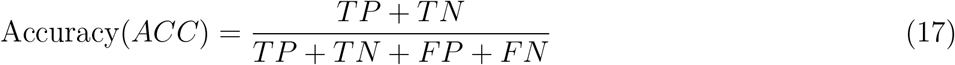

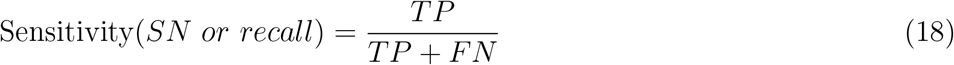

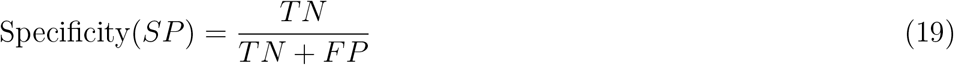

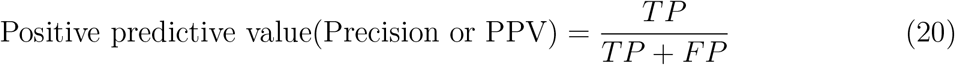

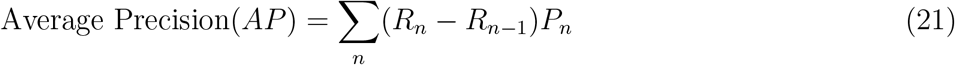

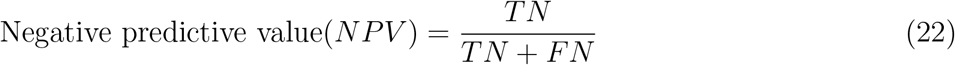

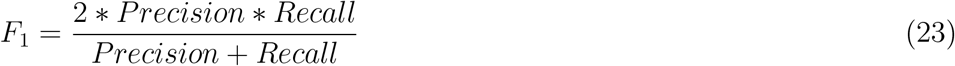

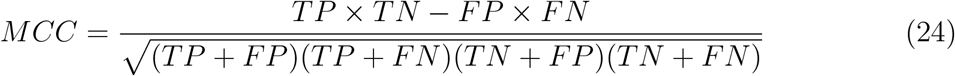

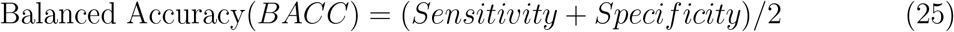

where *P*_*n*_ and *R*_*n*_ are the precision and recall at the *n*th threshold.

Given the highly imbalanced nature of the dataset (1:10 ratio of positives to negatives), traditional accuracy alone is insufficient to evaluate model performance. Instead, metrics such as F1-score, Matthews Correlation Coefficient (MCC), and Balanced Accuracy (BACC) were emphasized, as they provide more informative and class-sensitive assessments under skewed distributions. High sensitivity indicates effective detection of true interacting pairs, an elevated F1-score reflects a strong balance between precision and recall, MCC captures the overall quality of binary classification across all four confusion matrix components, and BACC ensures that both positive and negative classes are equally weighted by averaging sensitivity and specificity.

### 2.6 Resources

The ARACoFusion model was implemented using the PyTorch [26] deep learning framework for model construction and training. Evaluation metrics were computed using the scikit-learn library. Automated hyperparameter tuning was performed with Optuna. Model training and validation were conducted on a workstation equipped with Intel core i5 12400f and NVIDIA RTX 3060 GPU (12 GB VRAM).

## 3 Results

### 3.1 Performance comparison of protein language models

The performance of five different PLMs from ESM group and one from prot-T5 group [15] was compared (Figure3) before taking into consideration. Protein sequences were embedded using each of the PLM, the ARACoFusion model was trained on those embeddings with dataset C1 and tested on C2 and C3. In both the test datasets deeper transformer blocks consistently outperformed their shallower counterparts, with ESM-2 showing a clear incremental upgrade with increasing parameter size from T6 to T33. Despite similar parameter counts, the gated-ResNet architecture of ESM 1b [16] shows consistently increased performance compared to RoPE-enabled ESM2 [27] and Prot T5’s T5-styled decoder [15]. ESM-1b achieved the highest precision of 0.88 in both datasets C1 and C2. The obtained AUPR of ESM-1b were 0.86 on C2, 0.81 on C3, and the most pronounced margin can be witnessed in overall generalisation with MCC 0.73 over ProtT5 0.71. Collectively, these results confirm the robust predictive capabilities of deeper PLMs and highlight ESM 1b-T33 as the optimal choice for the protein-protein interaction task.

**Figure 3.**
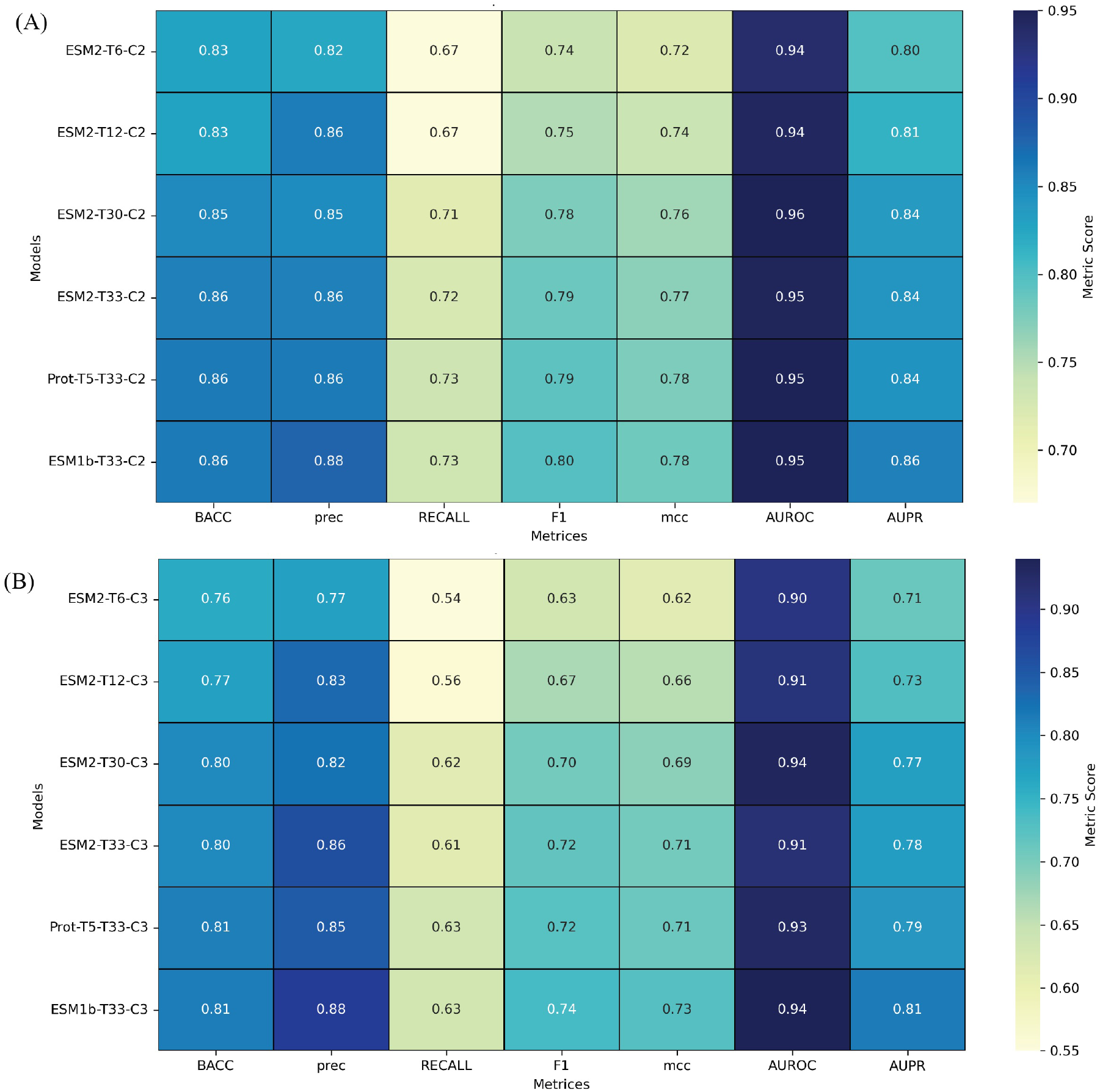
Comparison of pretrained protein language models on Arabidopsis PPI prediction. Heatmaps show model performance across (A) C2 and (B) C3 validation sets using BACC, Precision, Recall, F1, MCC, AUROC, and AUPRC.ESM-1b-T33 consistently achieves the highest AUPRC and F1, highlighting its superior generalization under class imbalance

### 3.2 Evaluation of Model Performance

#### 3.2.1 Effect of Hyperparameter Optimisation with Optuna

The Optuna framework was employed for Bayesian hyperparameter optimization to evaluate rigorously and maximize the predictive performance of ARACoFusion. We defined an objective function aimed at maximizing the area under the precision-recall curve (AUPRC) on the validation set. Optuna conducted automated optimization across a set of critical hyperparameters known to influence model training and generalization, including the number of attention heads in the reciprocal cross-attention encoder, learning rate for the AdamW optimizer, and Focal loss gamma (*γ*), dropout rate, label smoothing, and uncertainty weight (*λ*). The optimized configuration achieved superior performance compared to a baseline model trained with standard, untuned hyperparameters. The specific values for both configurations are summarized in Table2. To quantify the influence of each hyperparameter on model performance, we conducted a hyperparameter importance analysis using Optuna’s built-in ANOVA estimator. As illustrated in Figure4, the most impactful hyperparameter was the number of attention heads (importance = 0.30), highlighting the importance of cross-contextual modeling resolution. This was followed by learning rate (0.21) and focal loss gamma (0.16). Other parameters such as dropout, smoothing, and uncertainty weight also contributed, albeit to a lesser extent. To empirically validate the effectiveness of optimization, we conducted model evaluations on both the C2 and C3 datasets, comparing the Optuna-optimized configuration against the default baseline. The results (Tables3 and 4) demonstrate consistent improvements in recall, F1-score, MCC, and AUPRC, particularly in threshold-sensitive and imbalanced evaluation regimes. These findings confirm that targeted hyperparameter optimization plays a critical role in achieving robust generalization and calibrated decision-making in protein interaction prediction.

**Figure 4.**
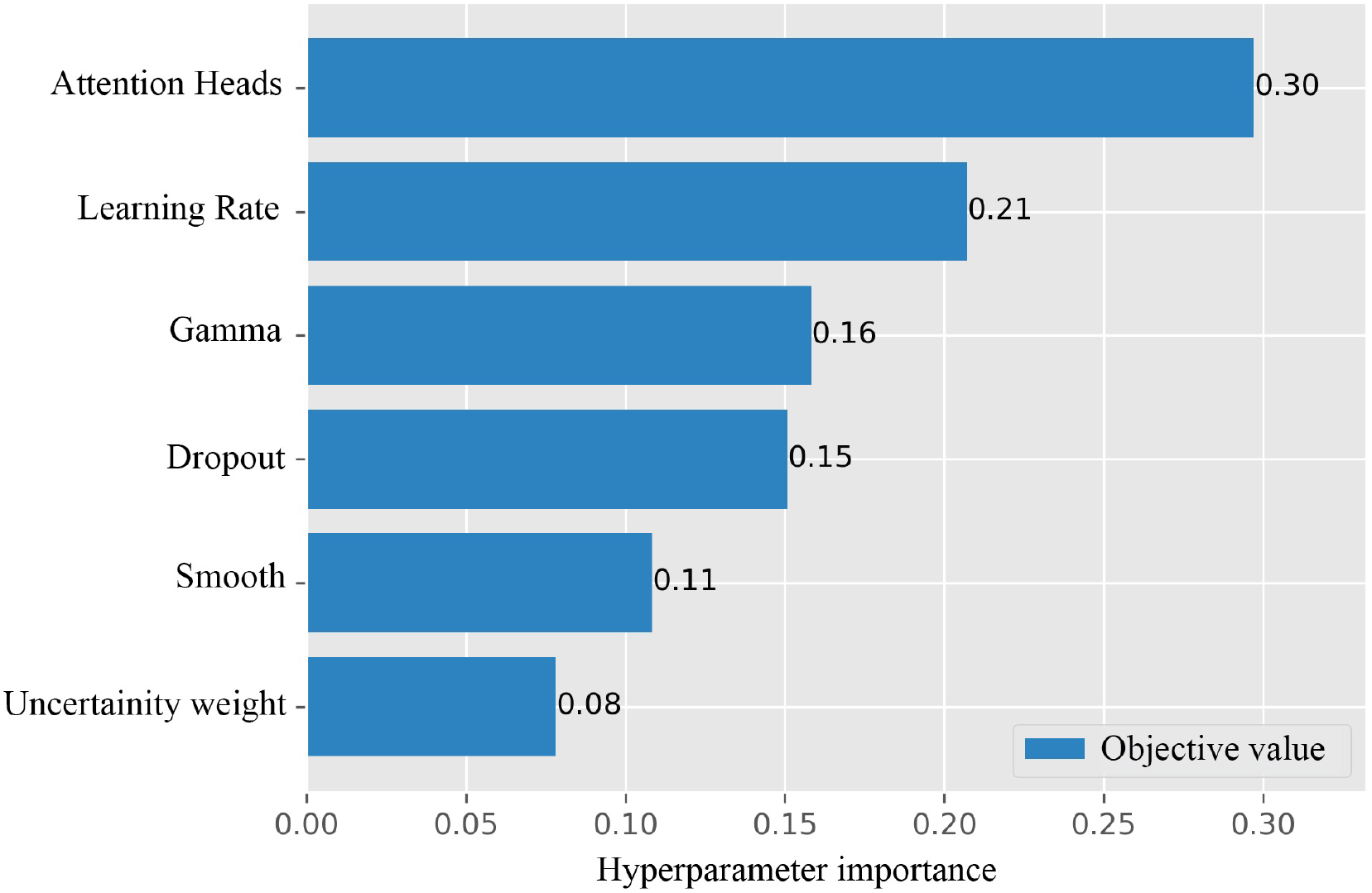
Hyperparameters importance graph obtained through Optuna

#### 3.2.2 tSne analysis

A two-dimensional t-SNE projection [28] represented (Figure 5) the raw ESM-1b-t33 pair embeddings for the imbalanced C2 split Figure 5(A) and the more compositionally diverse C3 split Figure 5(B) form largely overlapping manifolds in which interacting (red) and non-interacting (green) pairs are interspersed, defining the limited separability of sequence-only representations. In contrast, the ARACoFusion transformed embeddings for C2 (Figure 5(C)) and C3 (Figure 5(D)) exhibit pronounced class reorganization, interacting pairs as true positive indicated by green dots, consolidate into a dense core while non-interactors as true negative indicated by blue dots, migrate toward the periphery, yielding a clearer margin. Misclassified instances (false positives in yellow, false negatives in magenta) are confined mainly to the decision boundary, indicating that errors arise from borderline cases rather than systematic bias. The preservation of this topology across both splits implies that the ARACoFusion model learn interaction-relevant features that generalize beyond training distribution, complementing the quantitative gains reported in Table 2, Table 3, Table 4, and Table

**Table 2.** Best hyperparameters obtained through Optuna.

| Hyperparameters | Without Optuna | Best parameters with Optuna |
| --- | --- | --- |
| Attention Heads | 4 | 8 |
| Learning Rate | 0.001 | 0.000681383 |
| Gamma | 2.0 | 3.892573 |
| Dropout | 0.3 | 0.118479 |
| Smooth | 0.1 | 0.0514500 |
| Uncertainty weight | 0.1 | 0.579057 |

**Table 3.** ARACoFusion tested with and without Optuna on C2 data.

| DATASET C2 | ACC | SPEC | PREC | SN | F1 | MCC | NPV | AP | AUROC | BACC | AUPRC |
| --- | --- | --- | --- | --- | --- | --- | --- | --- | --- | --- | --- |
| No Optuna | **0.9658** | **0.9886** | **0.8659** | 0.738 | 0.7968 | 0.7812 | 0.9742 | 0.8451 | 0.9351 | 0.8633 | 0.8451 |
| With Optuna | 0.9657 | 0.9875 | 0.8568 | **0.7471** | **0.7982** | **0.7817** | **0.975** | **0.8546** | **0.9548** | **0.8673** | **0.8546** |

**Table 4.**
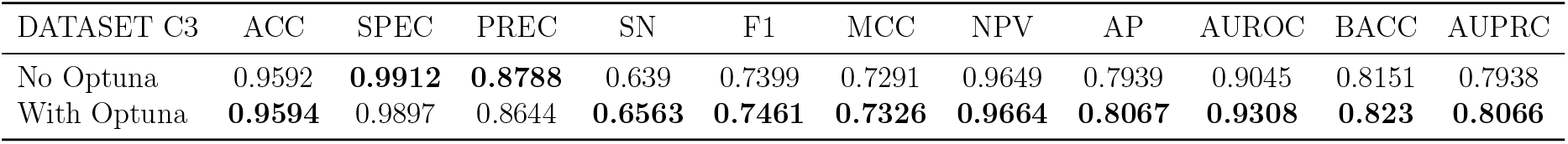
ARACoFusion tested with and without Optuna on C3 data.

**Figure 5.**
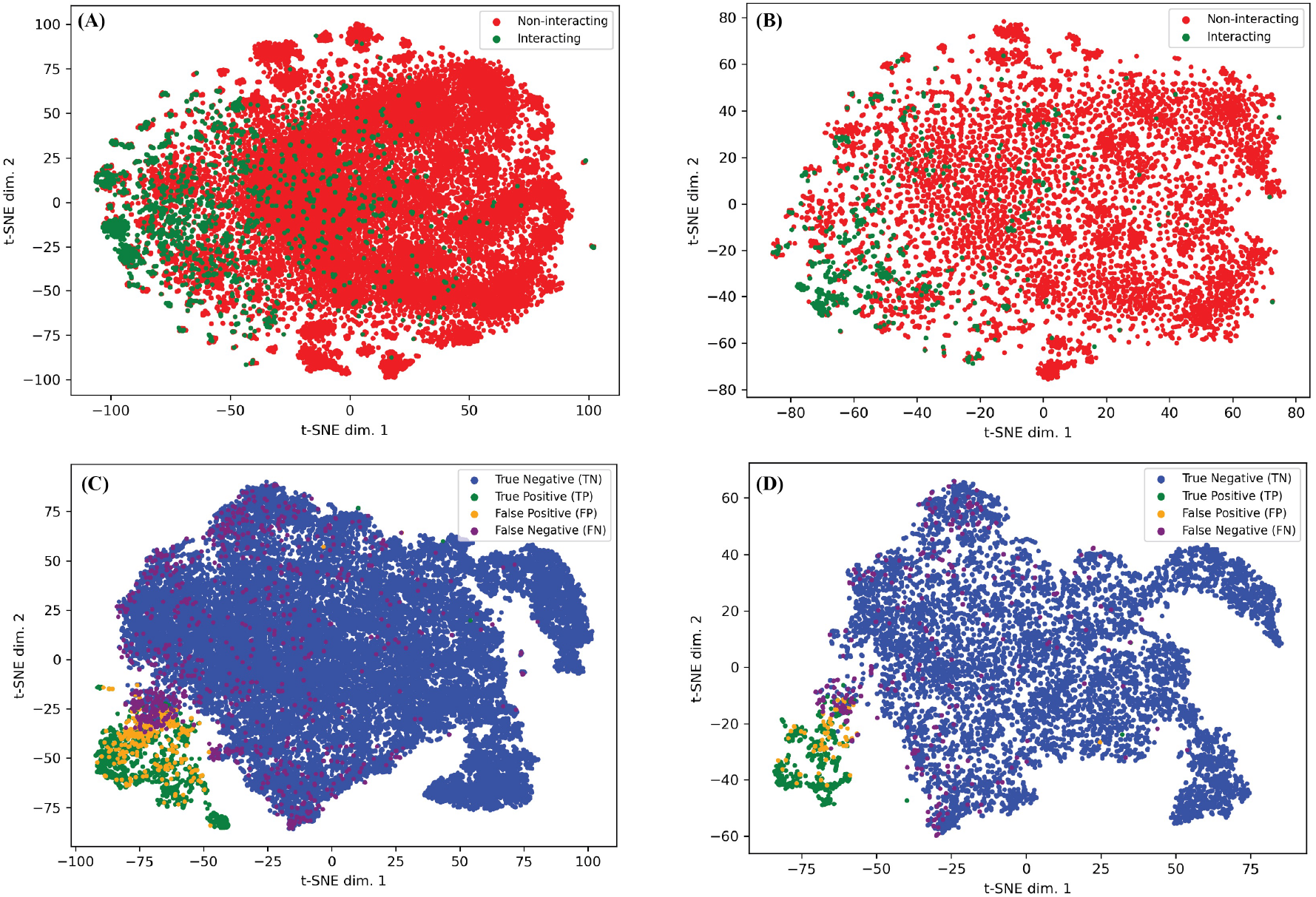
t-SNE plots comparing raw ESM-1b-t33 embeddings (A, B) with ARACoFusion-transformed embeddings (C, D) for C2 and C3 test sets. In (A, B), red and green indicate non-interacting and interacting pairs, respectively, showing overlapping distributions. In (C, D), transformed embeddings are colored by classification outcome: true negatives (blue), true positives (green), false positives (orange), and false negatives (magenta), revealing improved class separability and clearer decision boundaries after model transformation

#### 3.2.3 Calibration via temperature scaling

A reliability diagram assesses how well a model’s predicted probabilities align with actual observed outcomes; it puts predicted probabilities on the x-axis and the fraction of truly interacting pairs on the y-axis. If the model is accurately calibrated, every point falls on the 45° diagonal. The reliability diagram for the ARACoFusion classifier was illustrated (Figure 6), comparing raw prediction scores to scores recalibrated using temperature scaling (T). Specifically, temperature scaling adjusts the original logits by dividing each by an optimal constant T=0.58, determined on 10% test dataset to minimize the negative log-likelihood.The raw reliability curve (blue) exhibits non-uniform deviation from the identity line. It lies above the diagonal in certain lowand mid-confidence bins (around 0.3 to 0.4), indicating over-confidence, where the predicted probabilities are higher than the observed empirical frequencies. Conversely, in some regions (around 0.5 and 0.6), the curve dips below the diagonal, reflecting under-confidence. This inconsistency suggests that the model’s raw output lacks calibration, especially in the mid-probability range, where predicted scores do not reliably reflect the true likelihood of interaction. After applying temperature scaling with an optimal scalar T=0.58, the recalibrated curve (orange dashed) aligns more closely with the diagonal across all bins. This adjustment results in a notable improvement in calibration: the Expected Calibration Error (ECE) reduces from 0.034 (raw) to 0.020 (scaled), indicating that the predicted probabilities now better correspond to actual empirical interaction frequencies.

**Figure 6.**
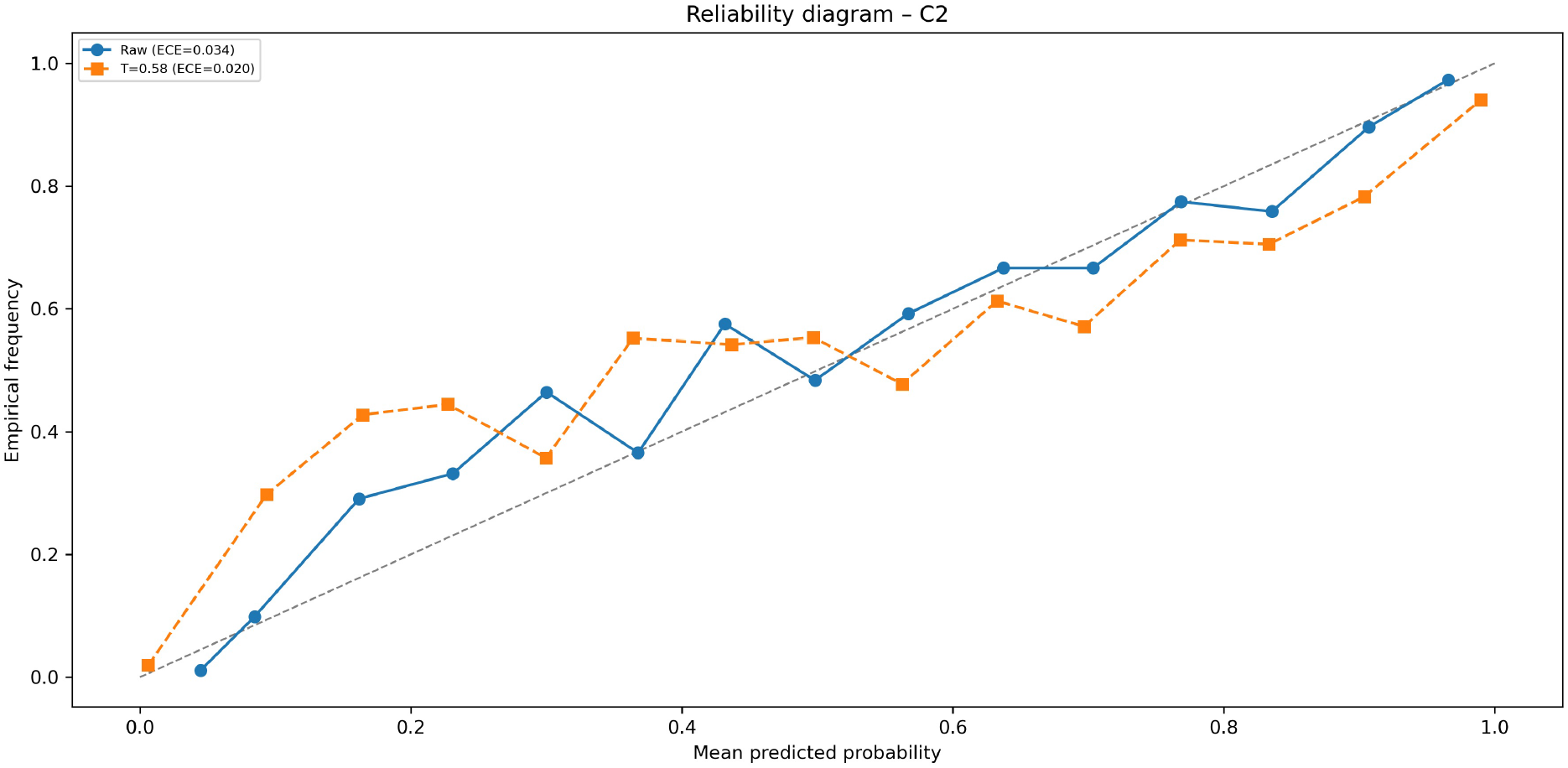
Reliability diagram for the C2 dataset before and after temperature scaling. The blue curve shows raw model outputs with an expected calibration error (ECE) of 0.034, while the orange curve represents calibrated predictions after applying temperature scaling (T = 0.58), reducing the ECE to 0.020.

#### 3.2.4 Benchmarking Against Existing Arabidopsis PPI Methods

The predictive efficacy of the proposed ARACoFusion architecture was evaluated comparing its performance against two established Arabidopsis protein-protein interaction (PPI) predictors: AraPPINet [19] and ESMAraPPI [21]. The results for ESMAraPPI were independently reproduced using the authors’ public GitHub repository, with original hyperparameter configurations preserved to ensure faithful replication. In contrast, due to missing implementation details for AraPPINet, its performance metrics were taken directly from the original publication. Evaluation was carried out under two complementary regimes: (i) split-based validation, wherein C1 was used as the training set and C2/C3 as test sets to probe generalization under class imbalance; and (ii) 5-fold cross-validation, performed on the merged C1+C2+C3 dataset to assess robustness under random partitioning. These benchmarking protocols reflect practical use cases in Arabidopsis interactome prediction, encompassing both class imbalance (due to scarce positive pairs) and distributional heterogeneity across data splits. Performance was assessed using a comprehensive suite of threshold-dependent metrics (e.g., F1-score, MCC, NPV) and threshold-independent criteria (AUROC, AUPRC).

#### 3.2.5 Comparing performance on split dataset

The results on the C2 dataset (Table 5) show that ARACoFusion significantly outperforms both ESMAraPPI and AraPPINet across nearly all evaluation metrics. It achieves an accuracy of 96.57%, with a substantially higher sensitivity 0.7471 and F1-score 0.7982 than its counterparts. Despite ESMAraPPI yielding a high precision of 0.9361, its sensitivity remains low 0.5038, indicating a conservative bias that suppresses detection of true positives. AraPPINet shows even lower sensitivity 0.337 and F1-score 0.551, despite extremely high specificity 0.999, thereby failing to recover a substantial portion of the interacting pairs.

**Table 5.** Comparison of ARACoFusion with existing Arabidopsis PPI predictors on C2 dataset.

| DATASET C2 | ACC | SPEC | PREC | SN | F1 | MCC | NPV | AP | AUROC | BACC | AUPRC |
| --- | --- | --- | --- | --- | --- | --- | --- | --- | --- | --- | --- |
| ARACoFusion | **0.9657** | 0.9875 | 0.8568 | **0.7471** | **0.7982** | **0.7817** | **0.975** | **0.8546** | **0.9548** | **0.8673** | **0.8546** |
| ESMAraPPI | 0.9518 | 0.9966 | 0.9361 | 0.5038 | 0.6551 | 0.6669 | 0.9526 | 0.8359 | **0.9665** | 0.7502 | 0.8358 |
| AraPPINet | 0.939 | **0.999** | **0.966** | 0.337 | N/A | 0.551 | N/A | N/A | N/A | N/A | N/A |

These findings are visually corroborated by Figure 7(A) and Figure 7(B), which displays the confusion matrices for ESMAraPPI and ARACoFusion on the C2 set. While both models correctly identify a large number of negatives (non-interacting pairs), ARACoFusion retrieves a notably higher number of true interacting pairs 2500 compared to ESMAraPPI only 1715, despite the severe class imbalance in which positives constitute a small minority. This demonstrates ARACoFusion’s enhanced recall capacity and resilience to imbalanced data.

**Figure 7.**
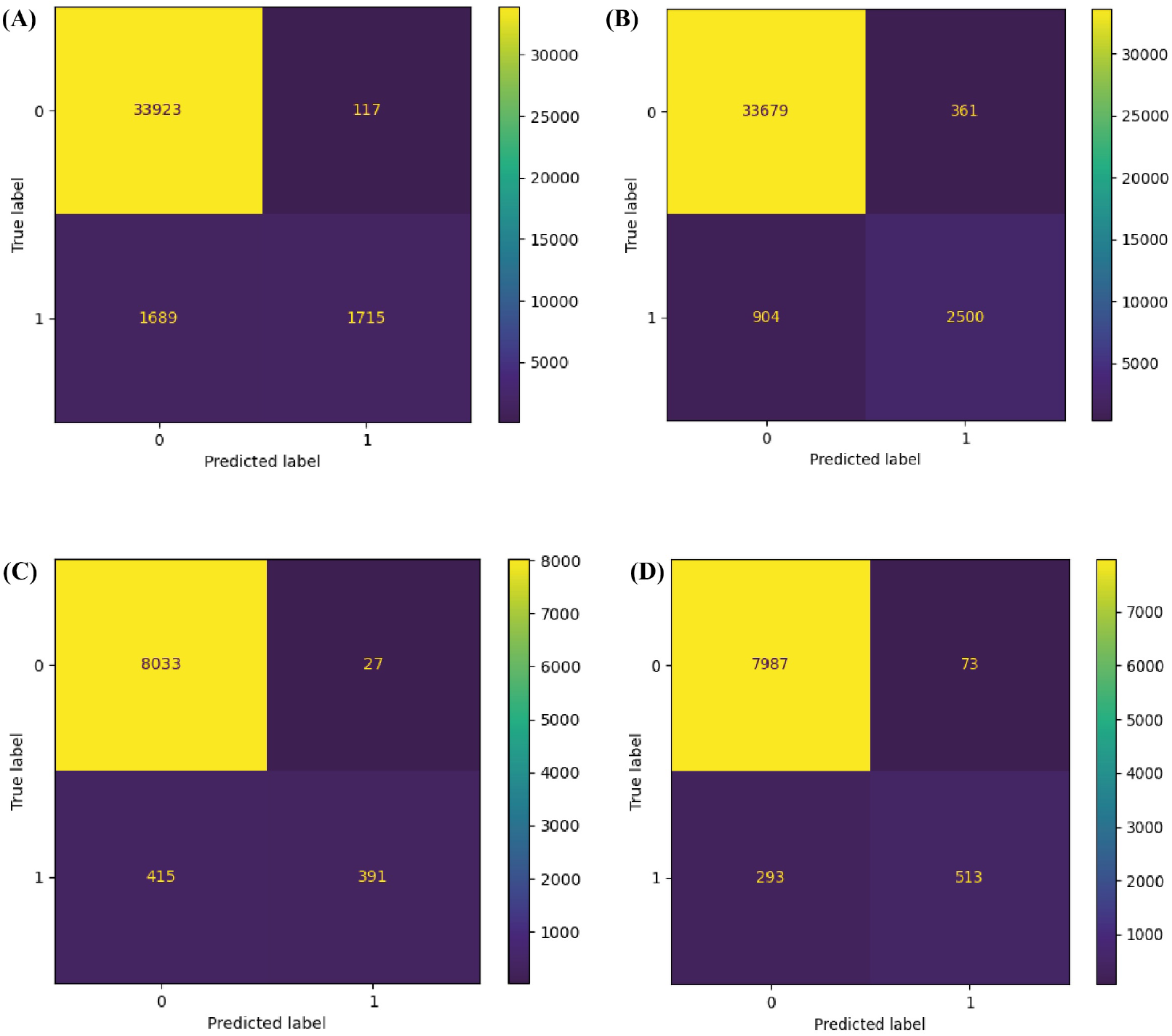
Comparing performance validation confusion matrices. (A) and (B) show performance on the C2 dataset for ESMAraPPI and ARACoFusion, respectively. (C) and (D) display the corresponding performance on the C3 dataset. Despite the significant class imbalance, ARACoFusion consistently recovers more true interactions, highlighting its superior sensitivity and robustness in low-prevalence contexts

On the C3 dataset, which introduces additional distributional variability, similar trends are observed (Table 7). ARACoFusion maintains a strong performance, achieving accuracy of 95.94%, sensitivity of 0.6563, and an AUPRC of 0.8066 with well above ES-MAraPPI’s sensitivity 0.4851 and AUPRC 0.8085. As depicted in Figure 6C and Figure 6D, ARACoFusion continues to recover more true positives 513 than ESMAraPPI 391, confirming its effectiveness in low-prevalence scenarios.

**Table 6.**
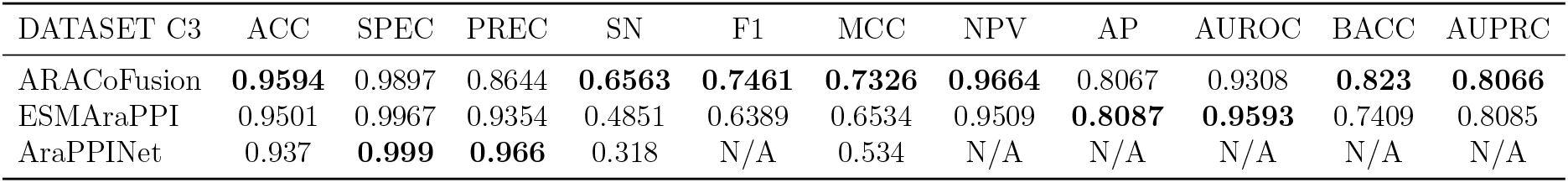
Comparison of ARACoFusion with existing Arabidopsis PPI predictors on C3 dataset.

**Table 7.**
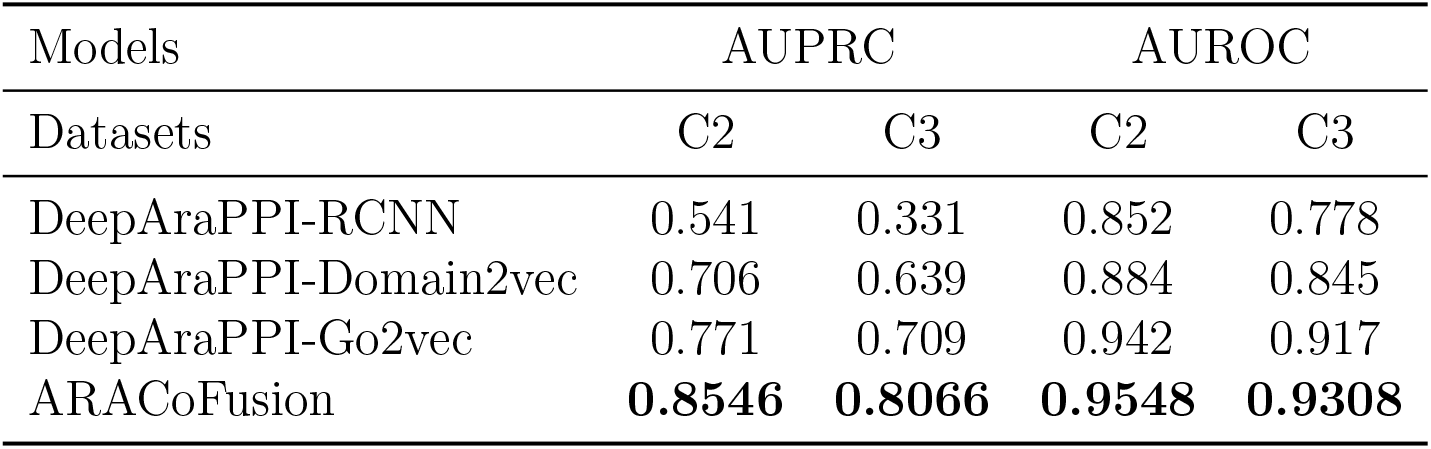
Comparing ARACoFusion with DeepAraPPI varients on C2 and C3 datasets.

In Table 5 we compared ARACoFusion with existing deep learning-based DeepAraPPI [20] variants across both splits, ARACoFusion outperforms all DeepAraPPI varients in AUPRC and AUROC, which are crucial for imbalanced classification tasks. Notably, on the C2 split, ARACoFusion achieves AUPRC = 0.8546, far exceeding DeepAra-PPI-RCNN 0.541 and GO2Vec 0.771. These margins persist in the C3 setting, reinforcing the generalizability of ARACoFusion.

#### 3.2.6 Comparing 5-fold cross-validation performance

A stratified 5-fold cross-validation was employed by merging the C1, C2, and C3 datasets into a unified Arabidopsis interactome benchmark to assess of the generalizability and robustness of the proposed model. This setup ensures that performance estimates are not biased by specific dataset compositions and provides a broader view of model behavior across varying interaction contexts. The average metrics over the 5 folds for both ARA-CoFusion and ESMAraPPI, the latter of which was evaluated using the original hyperparameters as provided in its public GitHub repository without further re−tuning (Table 8). This maintains consistency with the baseline architecture and prevents any form of implicit overfitting during re-evaluation. ARACoFusion achieves near-perfect generalization performance, with accuracy 99.78%, F1-score 0.988, and AUROC 0.9993, indicating extremely high fidelity across folds. Particularly notable is its sensitivity 0.9893, which again underscores its strength in correctly identifying positive interactions, even within class-imbalanced distributions. The model also maintains AUPRC 0.9967, demonstrating precision-recall stability under varied sampling partitions. In comparison, ESMAraPPI yields solid but consistently lower performance across all metrics. While the AUROC 0.9938 and AUPRC 0.9571 remain competitive, the drop in sensitivity 0.833 and F1-score 0.8799 highlights a recurring tradeoff: higher precision but lower recall. The drop in balanced accuracy to 91.36% further confirms that ESMAraPPI underperforms in recovering minority class instances (interacting pairs), echoing earlier trends seen in C2/C3 evaluations. These results reinforce the conclusion that ARACoFusion not only captures interaction-relevant features effectively but also generalizes reliably across diverse Arabidopsis PPI distributions.

**Table 8.** Comparing performance of ESMAraPPI with ARACoFusion with 5-fold cross validation.

| Models | ACC | SPEC | PREC | SN | F1 | MCC | NPV | AP | AUROC | BACC | AUPRC |
| --- | --- | --- | --- | --- | --- | --- | --- | --- | --- | --- | --- |
| ARACoFusion | **0.9978** | **0.9987** | **0.9867** | **0.9893** | **0.988** | **0.9868** | **0.9989** | **0.9967** | **0.9993** | **0.994** | **0.9967** |
| ESMaraPPI | 0.9795 | 0.9942 | 0.9359 | 0.833 | 0.8799 | 0.8714 | 0.9835 | 0.9571 | 0.9938 | 0.9136 | 0.9571 |

#### 3.2.7 Comparison to General PPI Prediction Methods

To contextualize the performance of ARACoFusion beyond Arabidopsis-specific baselines, we further compared its predictive efficacy against a range of general-purpose sequence-based PPI prediction methods. These include D-SCRIPT (Sledzieski et al., 2021), RAPPPID (Szymborski et al., 2022), PIPR (Chen et al., 2019), and TAGPPI (Song et al., 2022). All of which claim generalization across organisms. Evaluation was performed using the same C2 and C3 Arabidopsis test splits described previously, without additional fine-tuning of the baseline models. The comparative performance of the model is given in Table 9 of the metrics AUROC and AUPRC, the latter being especially critical under high imbalance. Across both C2 and C3 datasets, ARACoFusion outperforms all general models by substantial margins. On C2, it achieves an AUPRC of 0.8546, surpassing TAGPPI 0.700, the strongest general baseline. Similarly, for C3, ARACoFusion reaches an AUPRC of 0.8066, compared to TAGPPI’s 0.554. The gap is even more pronounced for earlier architectures like D-SCRIPT, which fails to cross an AUPRC of 0.30 on either split. AUROC shows ARACoFusion achieves 0.9548 (C2) and 0.9308 (C3), significantly ahead of TAGPPI (0.925 and 0.873, respectively). These results highlight the limitation of generic predictors when applied directly to plant-specific interactomes, and emphasize the effectiveness of training Arabidopsis-specialized architectures like ARACoFusion. Overall, this comparison underscores that species-adapted models with targeted sequence-attention architectures offer superior recall-precision tradeoffs, particularly in low-prevalence settings typical of plant interactomes.

**Table 9.** Comparing AUPRC and AUROC with general PPI prediction method.

| Models | AUPRC |  | AUROC |  |
| --- | --- | --- | --- | --- |
| Datasets | C2 | C3 | C2 | C3 |
| D-SCRIPT | 0.292 | 0.291 | 0.781 | 0.739 |
| RAPPPID | 0.516 | 0.371 | 0.852 | 0.800 |
| PIPR | 0.588 | 0.387 | 0.871 | 0.780 |
| TAGPPI | 0.700 | 0.554 | 0.925 | 0.873 |
| ARACoFusion | **0.8546** | **0.8066** | **0.9548** | **0.9308** |

#### 3.2.8 Cross-species generalization comparison

To assess the transferability of learned sequence representations across species boundaries, we evaluated the cross-species generalization capacity of ARACoFusion and ESMAraPPI, both trained exclusively on Arabidopsis C1 data, using an independent rice dataset for testing. The rice dataset presents an extreme class imbalance, with*∼*90% of pairs being non-interacting, thus simulating realistic low-prevalence scenarios in plant interactomics.

#### 3.2.9 Quantitative Evaluation

The cross-species performance of ARACoFusion and ESMAraPPI across 10 evaluation metrics is given in Table 10. Despite both models encountering a sharp decline in sensitivity, ARACoFusion achieves 0.0753 higher in recall than ESMAraPPI, while also obtaining 0.0581 higher AUPRC, a critical metric under high class imbalance. This indicates improved ability to retrieve true interacting pairs in a taxonomically divergent setting. The AUROC scores of ARACoFusion 0.017 lower than ESMAraPPI which is negligible and indicating our model is competitive, but AUROC tends to overestimate performance in imbalanced regimes and is thus less informative than AUPRC in this context.

**Table 10.** Comparing cross-species generalization performance.

| Models | ACC | SPEC | PREC | SN | F1 | MCC | NPV | AP | AUROC | BACC | AUPRC |
| --- | --- | --- | --- | --- | --- | --- | --- | --- | --- | --- | --- |
| ARACoFusion | 0.87 | 0.9195 | 0.3176 | **0.3748** | **0.3438** | **0.2734** | **0.9363** | 0.3525 | 0.6864 | **0.6471** | **0.3519** |
| ESMaraPPI | **0.8887** | **0.9476** | **0.3638** | 0.2995 | 0.3285 | 0.27 | 0.9312 | 0.2949 | **0.7034** | 0.6236 | 0.2938 |

These differences are visually reinforced in Figure 8, which presents confusion matrices for both models. ARACoFusion (Figure 8B) correctly identifies more true positives (220) than ESMAraPPI (Figure 8A) of 183, albeit at a slight cost to specificity. Importantly, ARACoFusion avoids over estimating negative values, capturing more positive interactions while maintaining acceptable false positive rates.

**Figure 8.**
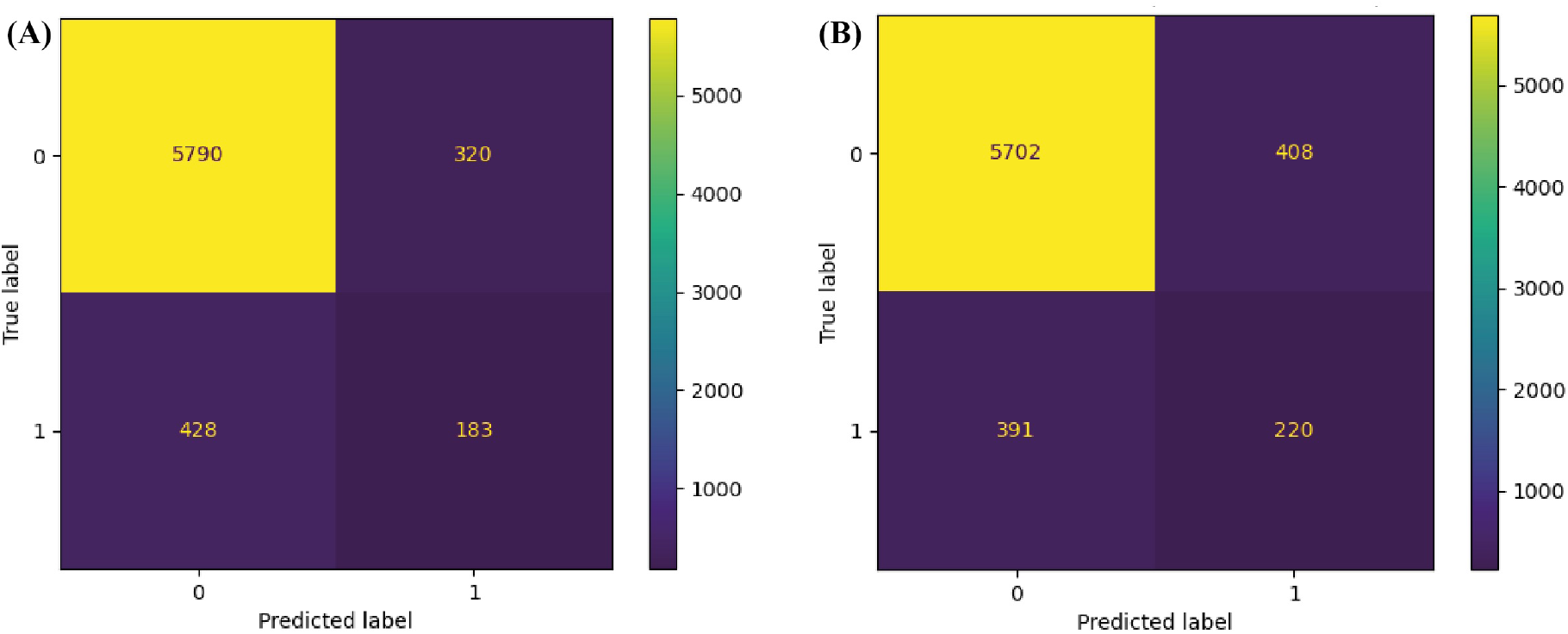
Comparison of model performance usning confusion matrix (A) performance of ESMAraPPI (B)Performance of ARACoFusion

#### 3.2.10 Embedding Quality via t-SNE Projection

To gain further insight into prediction accuarcy, we visualized t-SNE projections of rice sequence-pair embeddings before and after passing through ARACoFusion. As shown in Figure 9A, raw ESM-1b embeddings produce a dense, undifferentiated cluster with minimal visual separation between interacting (green) and non-interacting (red) pairs, reflecting the poor separability of pre-trained features in a cross-species context. In contrast, the ARACoFusion-transformed embeddings (Figure 9B) exhibit distinct substructures, where true positives (green) and true negatives (blue) occupy more cohesive, separable manifolds. False positives and false negatives are confined to the decision boundary, indicating that misclassifications arise primarily from borderline cases rather than systematic drift. These visual cues suggest that ARACoFusion learns to restructure the latent space in a way that reflects interacting and non-interacting proteins even on unseen species data.

**Figure 9.**
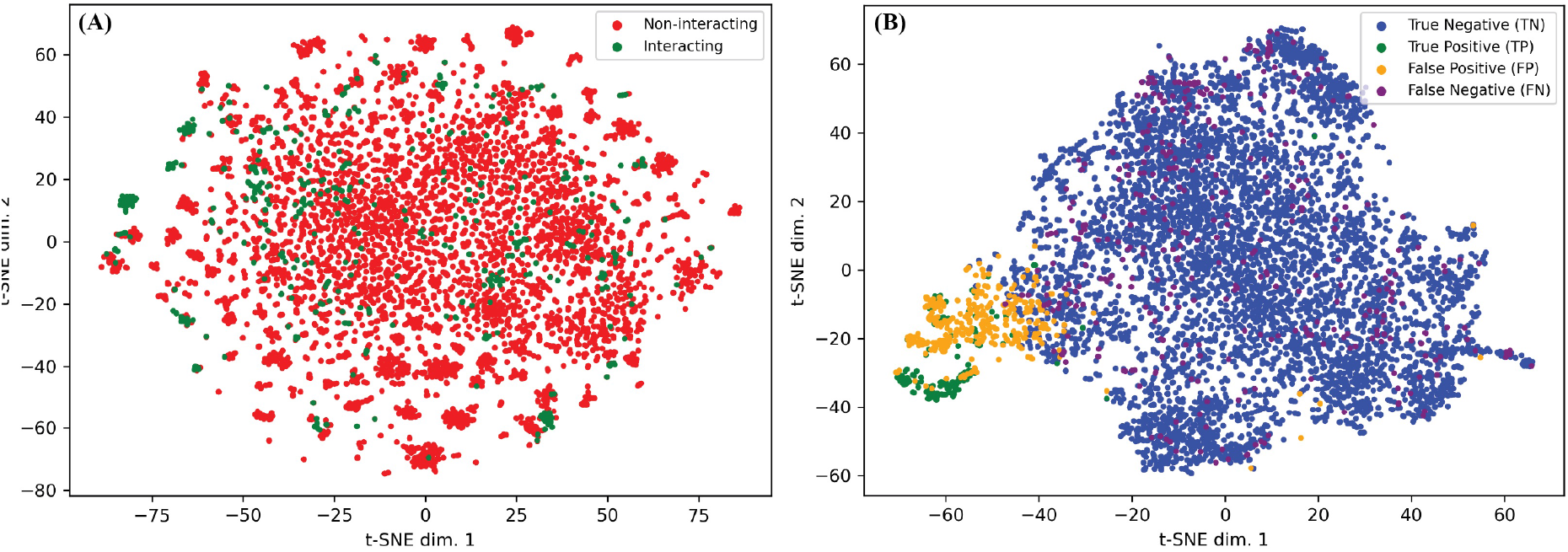
t-SNE projection of rice PPI embeddings. (A) Raw ESM-1b embeddings show poor separation between interacting (green) and non-interacting (red) pairs. (B) ARACoFusion transformed embeddings form distinct clusters, with misclassifications near the boundary, highlighting improved feature separability and cross-species generalization

#### 3.2.11 Comparison to Other Cross-Species Methods

To contextualize performance, we benchmarked ARACoFusion against existing cross-species or generic PPI classifiers, including RCNN (Chen et al., 2019), Domain2Vec, GO2Vec, Random Forest with DPC, and Logistic Regression (Zheng et al., 2023), as reported in Table 11. Across the board, ARACoFusion achieves the highest AUPRC (0.3519), outperforming GO2Vec (0.265) and Domain2Vec (0.279). Even traditional classifiers like LR (0.305) and RF+DPC (0.171) fall short of ARACoFusion’s precision-recall performance. These results collectively affirm that, while domain adaptation remains a challenge, sequence-informed attention models like ARACoFusion exhibit promising generalization to distantly related species. The model not only improves recall of rare true interactions but also restructures the decision space to better reflect interaction likelihood in taxonomically distinct proteomes.

**Table 11.** Cross-species AUPRC comparison of ARACoFusion with existing PPI classifiers.

| Datasets | AUPRC |
| --- | --- |
| RCNN | 0.248 |
| Domain2vec | 0.279 |
| GO2vec | 0.265 |
| RF+DPC | 0.171 |
| LR | 0.305 |
| ESMAraPPI | 0.2938 |
| ARACoFusion | **0.3519** |

#### 3.2.11 Evaluation on Arabidopsis Network Dataset

We performed a downstream evaluation on a biologically grounded interaction network derived from experimentally validated protein-protein interactions in Arabidopsis thaliana. It helped to further validate the practical applicability of ARACoFusion. The dataset was obtained from the STRING database [29] using stringent filtering criteria: (a) organism restricted to Arabidopsis thaliana, (b) interaction type limited to physical links only, (c) minimum confidence threshold set to 0.900, and (d) redundancy removed by retaining only AB (unidirectional) interactions. These filters yielded a high-confidence PPI network comprising 2477 proteins (nodes) and 9549 physical interactions (edges). The complete network was imported into Cytoscape (Shannon et al., 2025), and from it, we randomly selected a connected component consisting of 43 nodes and 103 edges for detailed case-study analysis. All edges in this subnetwork represent positive interactions according to STRING annotations. The prediction outcomes of the ARACoFusion model were illustrated in Figure 10 when applied to this isolated subnetwork. Edges colored in blue denote correctly predicted interactions, while red edges mark false positives or false negatives, depending on the binary classifier output. The green nodes represent the proteins involved; their STRING IDs were retained in the node. While ARACoFusion demonstrates an overall accurate reconstruction of the subnetwork, some false predictions persist, likely attributable to the negative-class bias in the training data (C1), which was generated with a large proportion of presumed non-interacting pairs. This introduces a degree of conservatism in classifying positive interactions, especially in densely connected modules. To benchmark the reliability of these predictions, we also tested the same subnetwork with ESMAraPPI, using identical pre-trained parameters. ARACoFusion achieved an accuracy of 70%, outperforming ESMAraPPI, which has 66% accuracy. These results highlight ARACoFusion’s improved alignment with experimentally derived PPI topologies and reinforce its utility in true biological network inference.

**Figure 10.**
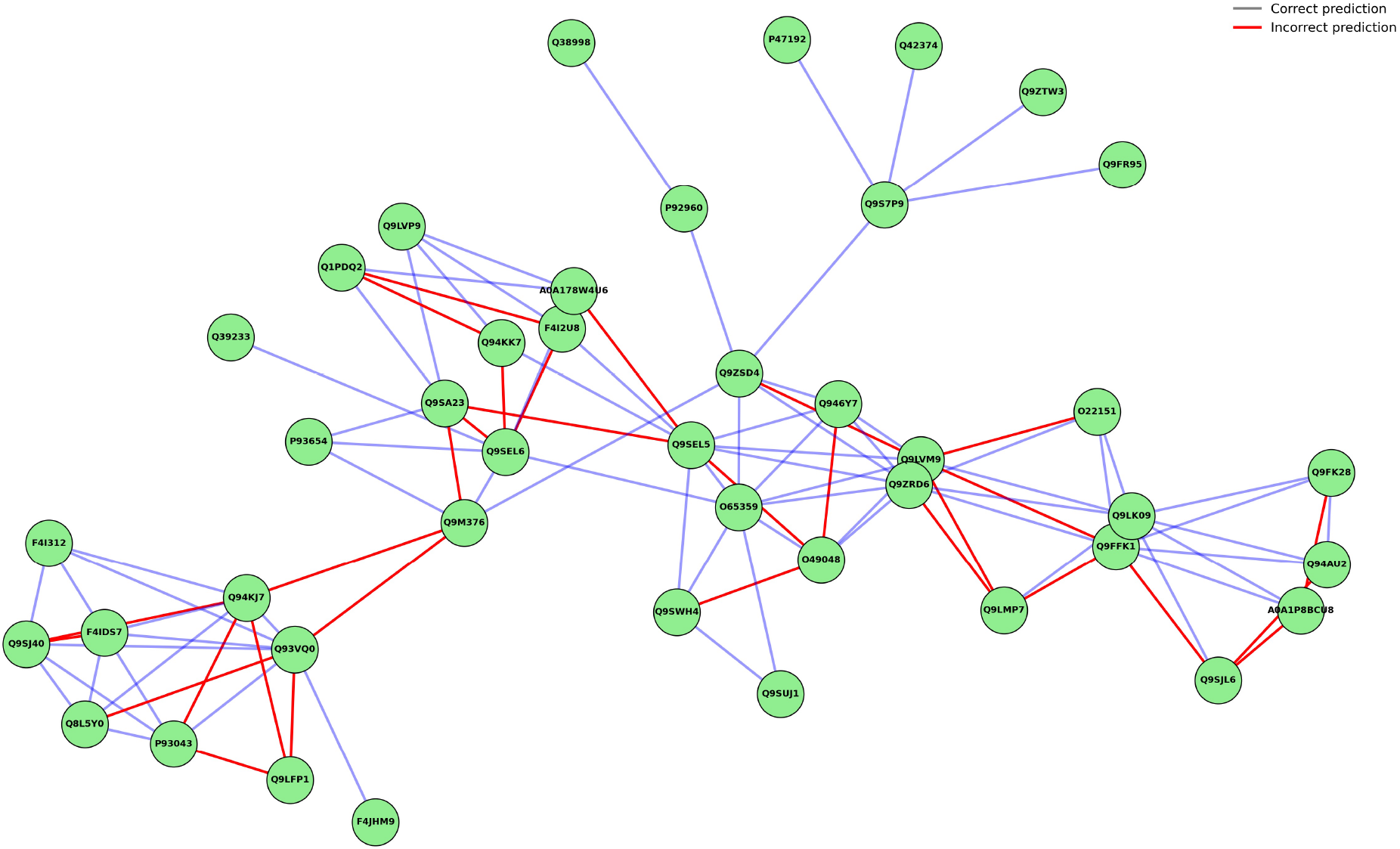
ARACoFusion prediction on a STRING-derived Arabidopsis PPI subnetwork. Blue edges indicate correct predictions; red edges show misclassifications. ARACoFusion achieved 70% accuracy, surpassing ESMAraPPI (66%), and effectively captured true interactions within a biologically validated network module

#### 3.2.13 Model Ablation Study

The individual contributions of key architectural and regularization components within ARACoFusion were assessed in ablation study. We conducted a systematic ablation study with four variant models were created by selectively removing one component at a time from the full architecture

a. the reciprocal cross-attention encoder,
b. the element-wise product and difference features,
c. the temperature scaling module, and
d. the uncertainty-aware training (variance regularization).

All ablation models were trained on the C1 dataset using identical training procedures and hyperparameters, and subsequently evaluated on the C2 and C3 test splits. This controlled setting isolates the impact of each component on the final performance.

##### Evaluation on C2 Dataset

The results on the C2 split (Table 12) indicate that the full ARACoFusion model delivers the highest sensitivity (0.7471), F1-score (0.7982), and balanced accuracy (0.8673). Removing the cross-attention encoder causes the largest performance drop in recall (down to 0.7051) and F1-score (0.787), suggesting that contextualized mutual interaction modeling is critical for positive interaction detection. Similarly, removing the element-wise product and absolute difference features slightly reduces precision and overall discriminability (MCC = 0.7686), affirming their role in capturing pairwise compatibility and divergence. When temperature scaling is removed, the model’s calibration capability is reduced, though the AUROC only shows a marginal drop (from 0.9559 to 0.9548). Interestingly, AUPRC increases slightly (0.8546 to 0.8555), suggesting that while calibration suffers, the precision-recall tradeoff remains stable or even marginally improved. Lastly, removing uncertainty-aware regularization has a subtle but consistent negative impact across nearly all metrics, reducing generalization and confidence calibration slightly.

**Table 12.** ARACoFusion ablation study on C2 dataset.

| Components removed from model | ACC | SPEC | PREC | RECALL | F1 | MCC | NPV | AP | AUROC | BACC | AUPRC |
| --- | --- | --- | --- | --- | --- | --- | --- | --- | --- | --- | --- |
| Reciprocal cross attention encoder | 0.9653 | 0.9913 | 0.8905 | 0.7051 | 0.787 | 0.7746 | 0.9711 | 0.8515 | 0.9524 | 0.8482 | 0.8515 |
| Element wise product and difference | 0.9637 | 0.9868 | 0.8474 | 0.7327 | 0.7859 | 0.7686 | 0.9736 | 0.8553 | 0.962 | 0.8597 | 0.8553 |
| temperature scaling | **0.966** | **0.9897** | **0.8759** | 0.7297 | 0.7962 | 0.7816 | 0.9734 | 0.8555 | 0.9559 | 0.8597 | 0.8555 |
| uncertainty aware training | 0.9647 | 0.9887 | 0.8655 | 0.7241 | 0.7885 | 0.7731 | 0.9729 | **0.8635** | **0.9693** | 0.8564 | **0.8635** |
| ARACoFusion (original) | 0.9657 | 0.9875 | 0.8568 | **0.7471** | **0.7982** | **0.7817** | **0.975** | 0.8546 | 0.9548 | **0.8673** | 0.8546 |

##### Evaluation on C3 Dataset

On the C3 dataset (Table 13), which is compositionally more diverse, similar trends are observed. The complete ARACoFusion model again achieves the highest sensitivity (0.6563) and MCC (0.7326), outperforming all ablated variants. Notably, the cross-attention removed variant yields the lowest sensitivity (0.6079) and F1-score (0.7275), indicating the centrality of inter-protein contextual encoding. The removal of temperature scaling and uncertainty regularization did not significantly harm performance in AUPRC (0.8138 and 0.8207, respectively, versus 0.8066 in the full model), though these components are known to enhance calibration and reliability rather than raw classification performance. Although the differences are not drastic in absolute accuracy, they become pronounced in threshold-sensitive metrics like MCC, AUPRC, and NPV, highlighting their utility in real-world, imbalanced prediction tasks. These findings collectively demonstrate that each component of the ARACoFusion architecture contributes synergistically to the final performance. The reciprocal crossattention encoder provides the most substantial boost by modeling inter-protein context. The latent interaction projector (element-wise operations) encodes explicit pairwise structure, enhancing compatibility encoding. Uncertainty-aware training improves confidence stability. Temperature scaling enhances post probability calibration without retraining. Thus, the ablation study confirms that ARACoFusion’s design is not an additive stack, but an interdependent system whose components jointly optimize generalization, robustness, and interpretability in sequence-based PPI prediction.

**Table 13.** ARACoFusion ablation study on C3 dataset.

| Components removed from model | ACC | SPEC | PREC | RECALL/SN | F1 | MCC | NPV | AP | AUROC | BACC | AUPRC |
| --- | --- | --- | --- | --- | --- | --- | --- | --- | --- | --- | --- |
| Reciprocal cross attention encoder | 0.9586 | **0.9937** | **0.9057** | 0.6079 | 0.7275 | 0.7225 | 0.962 | 0.8037 | 0.9326 | 0.8008 | 0.8036 |
| Element-wise product and difference | 0.9564 | 0.989 | 0.8509 | 0.6303 | 0.7242 | 0.7104 | 0.964 | 0.79 | 0.9376 | 0.8096 | 0.7899 |
| temperature scaling | 0.9589 | 0.9918 | 0.885 | 0.6303 | 0.7362 | 0.7268 | 0.9641 | 0.8139 | 0.9425 | 0.811 | 0.8138 |
| uncertainty aware training | 0.9577 | 0.9908 | 0.8722 | 0.6266 | 0.7292 | 0.7184 | 0.9637 | **0.8208** | **0.9582** | 0.8087 | **0.8207** |
| ARACoFusion (original) | **0.9594** | 0.9897 | 0.8644 | **0.6563** | **0.7461** | **0.7326** | **0.9664** | 0.8067 | 0.9308 | **0.823** | 0.8066 |

### 3.3 Webtool deployment

To facilitate broad accessibility and enable plant researchers to perform in silico PPI prediction without requiring computational infrastructure, we developed an interactive webtool implementing the ARACoFusion framework (Figure 11). The web interface accepts two protein sequences in and returns a calibrated interaction probability along with a confidence score derived from variance-aware estimation. Backend computations leverage precomputed ESM 1b-650M embeddings and execute the trained deep learning model using PyTorch and CUDA acceleration. The tool supports both single-pair predictions and batch submission for network-level inference. Additionally, users can visualize attention heatmaps and download structured prediction outputs for downstream integration. The webserver is hosted on a high-performance server infrastructure at https://ARAcofusion.compbiosysnbu.in/, ensuring rapid and stable performance. The documentation, and usage instructions are publicly available on GitHub.

**Figure 11.**
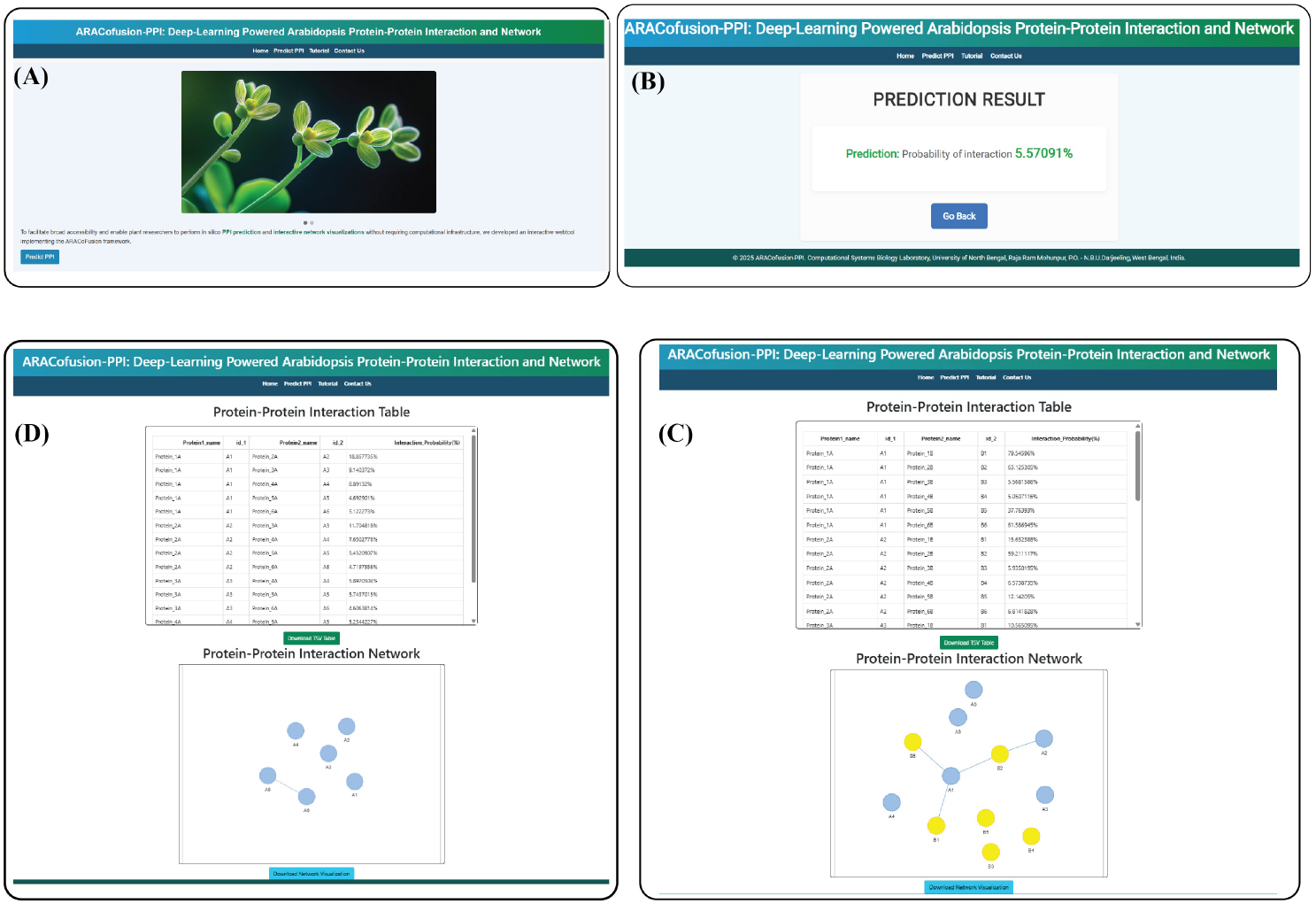
User interface of the ARACoFusion web server. The platform features: (A) A submission interface for entering raw protein sequences for analysis. (B) Results display showing the predicted interaction probability and confidence score for a specific pair. (C) Batch processing capabilities allowing users to upload datasets for group-based interaction screening. (D) Network visualization and prediction output for multi-sequence input files.

## 4 Conclusion

This study presents ARACoFusion, a domain-adapted deep learning framework that advances sequence-based protein-protein interaction prediction in plants. Building on the limitations of prior models, such as ESMAraPPI’s overconfident outputs and DeepAraPPI’s dependence on hand-crafted features. We designed a reciprocal attention-based architecture capable of modeling pairwise sequence relationships while maintaining scalability and interpretability. The use of pretrained ESM 1b 650M embeddings enables context-aware representation of primary protein sequences without requiring structural or evolutionary information, a critical advantage in plant systems where such annotations are rare. The reciprocal cross-attention encoder allows each protein in a pair to selectively attend to the other, capturing asymmetric binding cues and interaction-specific dependencies. This is further enhanced through latent projection and pairwise fusion operations (difference and product), which have been shown to improve downstream classification performance significantly during ablation. A major challenge addressed in this work is extreme class imbalance, with positive PPI instances comprising less than 10% of the dataset. To mitigate this, we employed focal loss with label smoothing, which reweights hard-to-classify positive samples while discouraging overfitting on large number of negatives. Moreover, we incorporated uncertainty-aware training by penalizing predictive variance across multiple forward passes, which led to more stable predictions and improved generalization. This is evident in our cross-species evaluation on the rice dataset, where ARACoFusion maintained strong AUPRC and MCC values, surpassing general-purpose PPI predictors. Importantly, we applied temperature scaling for post-prediction calibration of output probabilities, aligning predicted confidence scores with empirical correctness. This is critical for network-based applications where edge weights are interpreted probabilistically. Our calibration plots and reduced expected calibration error (ECE) demonstrate the practical utility of this step. Furthermore, hyperparameter optimization via Optuna contributed significantly to model tuning, offering a systematic alternative to manual grid or random search, ensuring model robustness across dataset partitions (C1, C2, C3). Benchmarking against state-of-the-art plant-specific models (DeepAraPPI, ESMAraPPI) and general PPI models confirmed the superiority of ARACoFusion across precision, recall, and balanced accuracy metrics. The t-SNE plots of learned representations illustrate clear class separation, while ablation results validate the contribution of each module. Finally, the deployment of ARACoFusionPPI as a webtool ensures usability by plant biologists in one-click-output, enabling large-scale, network-aware exploration of Arabidopsis interactomes. In summary, this work offers a calibrated, uncertainty-aware, and generalizable solution for plant PPI prediction, paving the way for future studies on network inference, context-specific PPI modeling, and integration with transcriptomic or phenotypic data.

## References

[1] Yixiang Zhang, Peng Gao, and Joshua S Yuan. Plant protein-protein interaction network and interactome. Current genomics, 11(1):40–46, 2010.

[2] Sebastian J Nintemann, Daniel Vik, Julia Svozil, Michael Bak, Katja Baerenfaller, Meike Burow, and Barbara A Halkier. Unravelling protein-protein interaction net-works linked to aliphatic and indole glucosinolate biosynthetic pathways in arabidopsis. Frontiers in plant science, 8:2028, 2017.

[3] Y Fang, D Macool, Z Xue, E Heppard, C Hainey, S Tingey, and G-H Miao. Development of a high-throughput yeast two-hybrid screening system to study protein-protein interactions in plants. Molecular Genetics and Genomics, 267(2):142–153, 2002.

[4] Jai S Rohila, Mei Chen, Shuo Chen, Johann Chen, Ronald L Cerny, Christopher Dardick, Patrick Canlas, Hiroaki Fujii, Michael Gribskov, Siddhartha Kanrar, et al. Protein-protein interactions of tandem affinity purified protein kinases from rice. PloS one, 4(8):e6685, 2009.

[5] Alisa G Woods, Izabela Sokolowska, Rama Yakubu, Melissa Butkiewicz, Martin LaFleur, Christopher Talbot, and Costel C Darie. Blue native page and mass spectrometry as an approach for the investigation of stable and transient protein-protein interactions. In Oxidative stress: diagnostics, prevention, and therapy, pages 341–367. ACS Publications, 2011.

[6] Yumeng Yan, D. Zhang, Pei Zhou, Botong Li, and Sheng-You Huang. Hdock: a web server for protein–protein and protein–dna/rna docking based on a hybrid strategy. Nucleic acids research, 45(W1):W365–W373, 2017.

[7] Takanori Hayashi, Yuri Matsuzaki, Keisuke Yanagisawa, Masahito Ohue, and Yutaka Akiyama. Megadock-web: an integrated database of high-throughput structure-based protein-protein interaction predictions. BMC bioinformatics, 19(Suppl 4):62, 2018.

[8] Jie Pan, Li-Ping Li, Zhu-Hong You, Chang-Qing Yu, Zhong-Hao Ren, and Yong-Jian Guan. Prediction of protein–protein interactions in arabidopsis, maize, and rice by combining deep neural network with discrete hilbert transform. Frontiers in Genetics, 12:745228, 2021.

[9] Yanzhi Guo, Lezheng Yu, Zhining Wen, and Menglong Li. Using support vector machine combined with auto covariance to predict protein–protein interactions from protein sequences. Nucleic acids research, 36(9):3025–3030, 2008.

[10] Bi-Qing Li, Kai-Yan Feng, Lei Chen, Tao Huang, and Yu-Dong Cai. Prediction of protein-protein interaction sites by random forest algorithm with mrmr and ifs. PLOS ONE, 7(8):1–10, 08 2012.

[11] Kyle H Ambert and Aaron M Cohen. K-information gain scaled nearest neighbors: a novel approach to classifying protein-protein interaction-related documents. IEEE/ACM Transactions on Computational Biology and Bioinformatics, 9(1):305– 310, 2011.

[12] Asif Ekbal, Sriparna Saha, Pushpak Bhattacharyya, et al. A deep learning architecture for protein-protein interaction article identification. In 2016 23rd international conference on pattern recognition (ICPR), pages 3128–3133. IEEE, 2016.

[13] Jie Pan, Zhu-Hong You, Li-Ping Li, Wen-Zhun Huang, Jian-Xin Guo, Chang-Qing Yu, Li-Ping Wang, and Zheng-Yang Zhao. Dwppi: a deep learning approach for predicting protein–protein interactions in plants based on multi-source information with a large-scale biological network. Frontiers in Bioengineering and Biotechnology, 10:807522, 2022.

[14] Ashish Vaswani, Noam Shazeer, Niki Parmar, Jakob Uszkoreit, Llion Jones, Aidan N Gomez, Lukasz Kaiser, and Illia Polosukhin. Attention is all you need. Advances in neural information processing systems, 30, 2017.

[15] Ahmed Elnaggar, Michael Heinzinger, Christian Dallago, Ghalia Rehawi, Yu Wang, Llion Jones, Tom Gibbs, Tamas Feher, Christoph Angerer, Martin Steinegger, et al. Prottrans: Toward understanding the language of life through self-supervised learning. IEEE transactions on pattern analysis and machine intelligence, 44(10):7112– 7127, 2021.

[16] Alexander Rives, Joshua Meier, Tom Sercu, Siddharth Goyal, Zeming Lin, Jason Liu, Demi Guo, Myle Ott, C Lawrence Zitnick, Jerry Ma, et al. Biological structure and function emerge from scaling unsupervised learning to 250 million protein sequences. Proceedings of the National Academy of Sciences, 118(15):e2016239118, 2021.

[17] Nadav Brandes, Dan Ofer, Yam Peleg, Nadav Rappoport, and Michal Linial. Proteinbert: a universal deep-learning model of protein sequence and function. Bioinformatics, 38(8):2102–2110, 2022.

[18] Dipayan Sarkar and Chiranjib Sarkar. Attnseq-ppi: Enhancing protein-protein interaction network prediction using transfer learning-driven hybrid attention. Biochimica et Biophysica Acta (BBA)-Proteins and Proteomics, page 141102, 2025.

[19] Jiawei Zhao, Yu Lei, Jianwei Hong, Cunjian Zheng, and Lida Zhang. Arappinet: an updated interactome for the analysis of hormone signaling crosstalk in arabidopsis thaliana. Frontiers in Plant Science, 10:870, 2019.

[20] Jingyan Zheng, Xiaodi Yang, Yan Huang, Shiping Yang, Stefan Wuchty, and Ziding Zhang. Deep learning-assisted prediction of protein–protein interactions in arabidopsis thaliana. The Plant Journal, 114(4):984–994, 2023.

[21] Kewei Zhou, Chenping Lei, Jingyan Zheng, Yan Huang, and Ziding Zhang. Pretrained protein language model sheds new light on the prediction of arabidopsis protein–protein interactions. Plant Methods, 19(1):141, 2023.

[22] Takuya Akiba, Shotaro Sano, Toshihiko Yanase, Takeru Ohta, and Masanori Koyama. Optuna: A next-generation hyperparameter optimization framework. In Proceedings of the 25th ACM SIGKDD international conference on knowledge discovery & data mining, pages 2623–2631, 2019.

[23] Yungki Park and Edward M Marcotte. Flaws in evaluation schemes for pair-input computational predictions. Nature methods, 9(12):1134–1136, 2012.

[24] Yarin Gal and Zoubin Ghahramani. Dropout as a bayesian approximation: Representing model uncertainty in deep learning. In international conference on machine learning, pages 1050–1059. PMLR, 2016.

[25] Chuan Guo, Geoff Pleiss, Yu Sun, and Kilian Q Weinberger. On calibration of modern neural networks. In International conference on machine learning, pages 1321–1330. PMLR, 2017.

[26] Adam Paszke, Sam Gross, Francisco Massa, Adam Lerer, James Bradbury, Gregory Chanan, Trevor Killeen, Zeming Lin, Natalia Gimelshein, Luca Antiga, et al. Pytorch: An imperative style, high-performance deep learning library. Advances in neural information processing systems, 32, 2019.

[27] Zeming Lin, Halil Akin, Roshan Rao, Brian Hie, Zhongkai Zhu, Wenting Lu, Nikita Smetanin, Robert Verkuil, Ori Kabeli, Yaniv Shmueli, et al. Evolutionary-scale prediction of atomic-level protein structure with a language model. Science, 379(6637):1123–1130, 2023.

[28] Laurens van der Maaten and Geoffrey Hinton. Visualizing data using t-sne. Journal of machine learning research, 9(Nov):2579–2605, 2008.

[29] Damian Szklarczyk, Rebecca Kirsch, Mikaela Koutrouli, Katerina Nastou, Farrokh Mehryary, Radja Hachilif, Annika L Gable, Tao Fang, Nadezhda T Doncheva, Sampo Pyysalo, et al. The string database in 2023: protein–protein association networks and functional enrichment analyses for any sequenced genome of interest. Nucleic acids research, 51(D1):D638–D646, 2023.

